# Resolving noise-control conflict by gene duplication

**DOI:** 10.1101/634741

**Authors:** Michal Chapal, Sefi Mintzer, Sagie Brodsky, Miri Carmi, Naama Barkai

## Abstract

Gene duplication promotes adaptive evolution in two principle ways: allowing one duplicate to evolve a new function and resolving adaptive conflicts by splitting ancestral functions between the duplicates. In an apparent departure from both scenarios, low-expressing transcription factor (TF) duplicates commonly regulate similar sets of genes and act in overlapping conditions. To examine for possible benefits of such apparently redundant duplicates, we examined the budding yeast duplicated stress regulators Msn2 and Msn4. We show that Msn2,4 indeed function as one unit, inducing the same set of target genes in overlapping conditions, yet this two-factor composition allows its expression to be both environmental-responsive and with low-noise, thereby resolving an adaptive conflict that inherently limits expression of single genes. Our study exemplified a new model for evolution by gene duplication whereby duplicates provide adaptive benefit through cooperation, rather than functional divergence: attaining two-factor dynamics with beneficial properties that cannot be achieved by a single gene.

## Introduction

The number of Transcription Factors (TFs) expressed in eukaryotes increases rapidly with increasing genome size and organism complexity, ranging from ∼50 in obligate parasites to >1000 in high eukaryotes^1^. Gene duplication played a major role in this evolutionary expansion^2,3^, as is evident from the fact that the number of DNA Binding Domains (DBD) remained practically constant with the increasing genome size. In fact, the majority of TFs belong to just a few DBD families^2,3^, the content of which increased rapidly with genome size. Understanding the adaptive forces that promote this duplication-dependent expansion is therefore of a great interest.

Gene duplication can promote evolution by allowing one of the duplicates to adapt a novel function while the second duplicate maintains the ancestral function. More often, however, the two duplicates do not gain a new function but rather lose complementary subsets of ancestral functions^4,5^. Sub-functionalization not only explains duplicate maintenance, but can also promote adaptive evolution by enabling further optimization of each individual function and resolving adaptive conflicts of the ancestral gene^6,7^. Indeed, optimizing a dual-function protein is often constrained by conflicting requirements imposed by the different functions: a mutation that favor one function can perturb the other function, presenting an adaptive conflict that is only resolved upon duplication.

In the context of TFs, duplication may allow one factor to acquire a new set of target genes (neo-functionalization). Alternatively, the ancestral targets could split between the duplicates (sub-functionalization). In both scenarios, duplicate divergence would increase and refine the regulatory logic. Previous studies exemplified both scenarios^8–10^, but whether they are relevant for the majority of TF duplicates remained unclear. In fact, present-day genomes express a large number of TF duplicates that regulate highly similar, or even redundant, set of targets.

Budding yeast provide a convenient platform for studying the possible adaptive role of apparently redundant duplicates. The yeast lineage underwent a Whole Genome Duplication (WGD) event about one hundred million years ago^11^, and while most duplicates generated in this event were lost, about 10% were retained, amongst which TF are over-represented. Many of the retained TF duplicates show little signs of divergence but rather retained a highly conserved DBD, and, accordingly, bind the same DNA binding motif and regulate a highly similar set of genes. We reasoned that studying such duplicates might help shed light on possible benefits provided by TF duplication.

As a case in point, we investigated Msn2 and Msn4, a well-studied TF duplicates in budding yeast, which activate a large number of targets of the environmental stress response^12,13^. Previous studies established the tight similarity between these factors^14^, but we still began our study by increasing experimental resolution in an attempt to detect target divergence. Our results, however, re-enforced the conclusion that the two factors regulate the same set of target genes, translocate to the nucleus with the precise same dynamics, and expressed in an overlapping set of conditions.

Our search for differences between the duplicates pointed us in a different direction: the challenge which cells face when attempting to minimize noise in gene expression. Transcription is a stochastic process and is therefore characterized by random variations (noise) between genetically identical cells^15,16^. Cell to cell expression variability is deleterious for genes that require precise tuning of expression, such as dosage sensitive genes^17,18^, but beneficial when enabling processes not possible by deterministic dynamics, such as bet-hedging strategies^19–21^. Accordingly, noise levels vary greatly between genes^22^. Yet, the ability to tune expression noise through changes in gene promoter is limited by mechanistic constraints, and in particular by the well-documented conflict between regulatory control and noise: genes that are regulated over a wide dynamic range also show a high level of expression noise^15,23–25^. Thus, while coding for low-noise expression is possible, it comes at the cost of lowering the dynamic range over which expression can be changed by regulatory signals.

Our study shows that through duplication, Msn2,4 resolved this interplay between environmental responsive and noise. Following duplication, Msn2 expression became highly stable, showing limited responsiveness to environmental conditions and low expression noise. By contrast, Msn4 expression accentuated the environmental-responsive expression of the unduplicated homologue. This resulted in an overall expression of the Msn2,4 unit that is responsive to the environment but is also low-noise at the low expressing conditions. We provide evidence that this expression tuning is phenotypically adaptive, and define the genetic changes that correlates with the change in gene responsiveness and noise. Our results suggest that duplicates can promote adaptive evolution not only through functional divergence, as suggested by the neo-or sub-functionalization models, but also through effective cooperation, by attaining two-factor dynamics with emergence beneficial properties that cannot be achieved using a single gene.

## Results

### Low-noise (Poisson) distribution of MSN2 expression in individual cells

Msn2 and Msn4 are TF duplicates that regulate the stress response in budding yeast^12,26^. Stress genes show a noisy expression, and we were therefore surprised to observe that Msn2 is expressed at very similar amounts across individual cells. In fact, Msn2-GFP was the least noisy of all proteins expressed at its level, as quantified in a study surveying >2500 GFP-fused proteins^22^ (Figure 1A). To examine whether this low noise is also seen at the *MSN2* transcript level, we used single-molecule Fluorescent In-Situ Hybridization (smFISH)^27^ technique (Figure 1B). The number of *MSN2* transcripts in individual cells was well described by a Poisson distribution, as expected when individual mRNA transcripts are produced and degraded at constant rates^28,29^. This distribution presents the lower limit of gene expression noise, obtained in the absence of regulation, and other noise-amplifying processes^28^.

**Figure 1.**
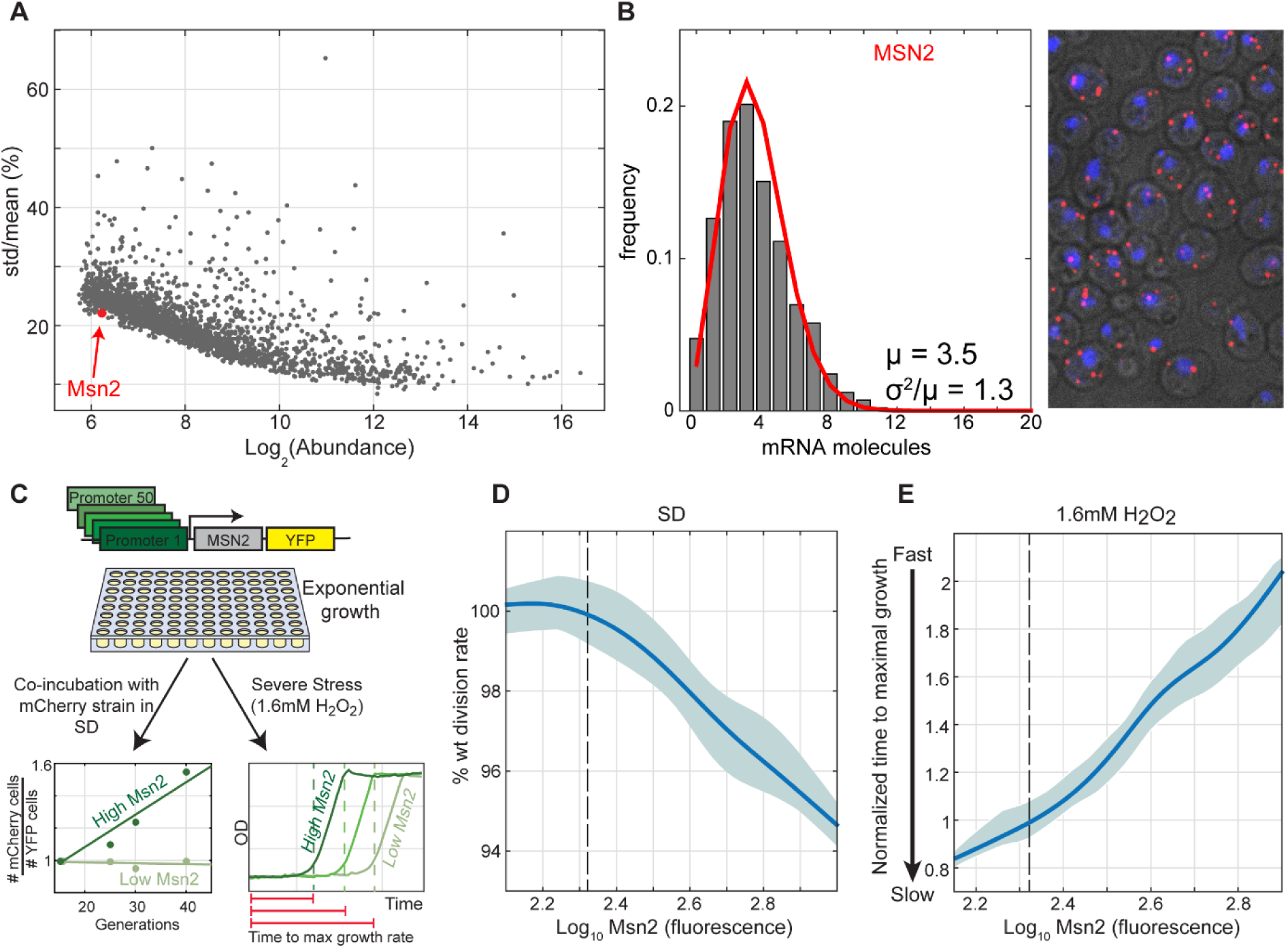
Tuning of Msn2 expression in rapidly growing cells: **(A)** *Cell-to-cell variability of Msn2-GFP is the lowest of all equally abundant proteins:* Shown is the noise vs. abundance data of ∼2,500 GFP-fused proteins, data from: Newman et al.^22^ Msn2 is shown as a red dot. Msn4-GFP was not detected. **(B)** *Low-noise (Poisson) distribution of MSN2 in individual cells: MSN2* expression levels were measured using smFISH. (Left) *MSN2* mRNA counts distribution, quantified in >650 single cells. Red line represents Poisson fit to the data. (Right) Fixed cells labeled with *MSN2* mRNA in red and DAPI staining in blue, in a maximal z-projection image. (**C-E**) *Msn2 expression increase stress protection by compensate growth rate in the absence of stress*: A library of 50 strains expressing Msn2-YFP under different synthetic promoters (from Keren et al.^33^) defining a range of expression values was generated (**C,** Methods). This library was used to define the effect of Msn2 expression level on growth rate and stress protection. Growth rates were measured using a sensitive competition assay, and is shown in (**D).** Stress protection was measured by subjecting exponentially growing cells to H_2_O_2_ (1.6mM), and identifying the time at which growth was first detected by continuous OD measurements (**E).** Shown are the median of all strains and repeats in solid line and 25-75 percentiles in the shaded areas. Dashed lines indicate wild-type Msn2 level.

### Increasing Msn2 expression promotes stress protection but reduces cell growth rate

Low expression noise characterizes genes coding for essential functions or components of large complexes^29,30^, for which expression tuning is beneficial^30–32^. By contrast, Msn2 is not essential, does not participate in large complexes, and is mostly inactive in rich media. To examine whether, and how, Msn2 expression level impacts cell fitness, we engineered a library of strains expressing Msn2 at gradually increasing amounts using synthetic promoters^33^. Measuring growth rates of the library strains using a sensitive competition assay (Figure 1C), we found that decreasing Msn2 expression to below its wild-type levels, and down to a complete deletion, had no detectable effect on growth rate within the resolution of our assay (0.5%). By contrast, increasing Msn2 abundance gradually decreased growth rate (Figure 1D,S1). Next, we tested the effect of Msn2 levels on the ability to proliferate in harsh stress, by incubating the library cells with high H_2_O_2_ concentrations (Figure 1C,E,S1). Here, increasing Msn2 levels was beneficial: cells that expressed high levels of Msn2 resumed growth faster than low-expressing ones. Therefore, increasing Msn2 expression better protects cells against stress, but reduces their growth rate. An optimal Msn2 level is therefore desirable to balance the need for rapid growth and stress protection, explaining the requirement for low-noise tuning of its gene expression.

The tradeoff between rapid growth and stress preparation depends on the contribution of these two parameters to the overall population fitness, as defined by the evolutionary history. This relative contribution, in turn, depends on growth conditions. For example, when cells encounter optimal growth conditions, maximizing division rate dominates, but when nutrients become limiting, or respiration is triggered, the importance of stress protection increases. Consistent with this, wild-type cells were better protected against H_2_O_2_ exposure at higher cell densities, as they approached stationary phase, resuming growth faster after stress induction (Figure 2A). We therefore expected Msn2 expression to increase with cell density. This, however, was not the case. Although Msn2 contributed to stress protection at all densities, its expression remained constant, throughout the growth curve (Figure 2B).

**Figure. 2.**
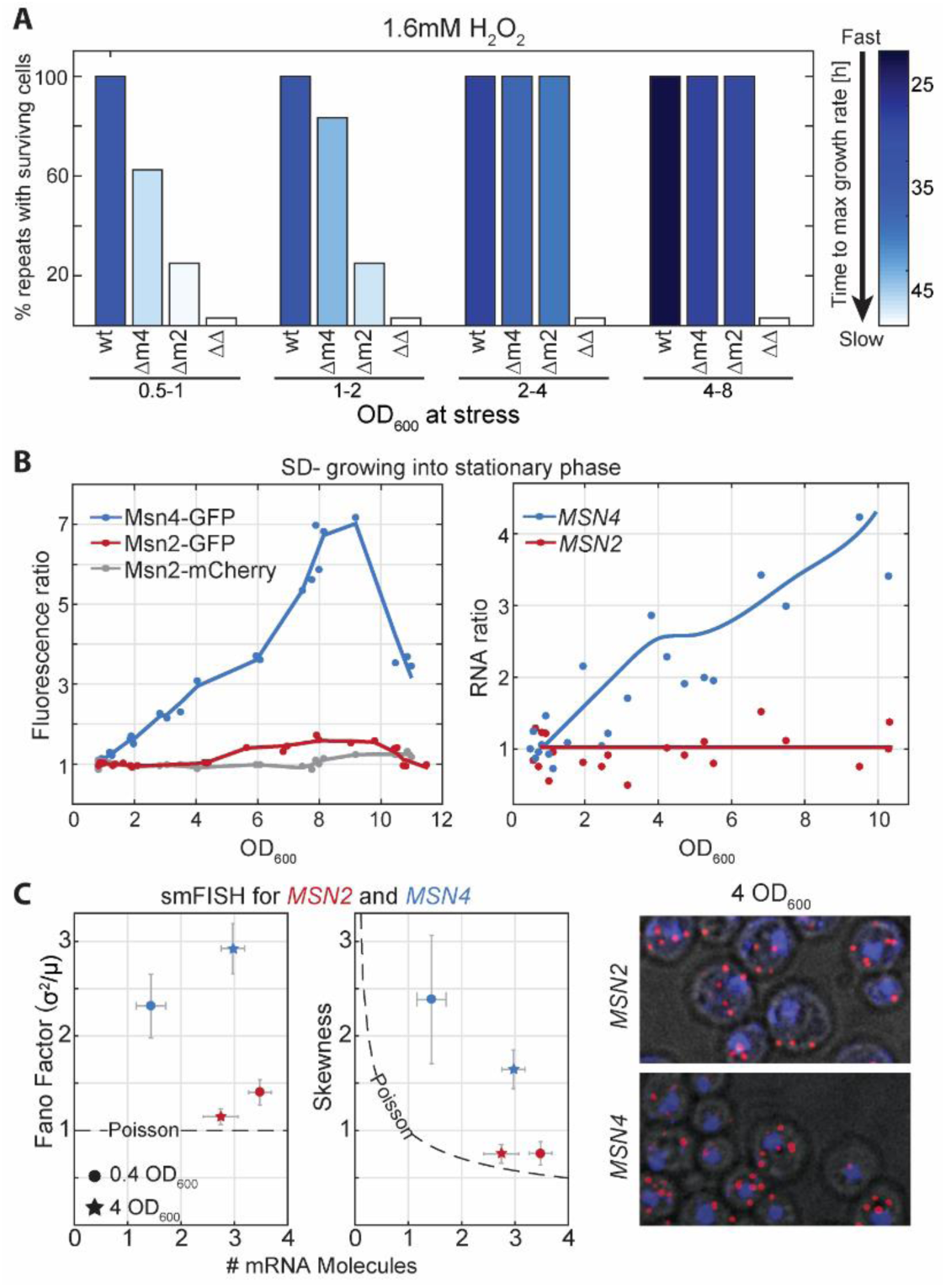
Msn4 expression and its contribution to stress preparation increases as cells exit exponential growth: **(A)** *The contribution of Msn2 and Msn4 to stress preparation changes along the growth curve:* Cells at different stages along the growth curve (ODs) were diluted into media containing 1.6mM H_2_O_2_ and were followed by continuous OD measurements to define the time at which growth was first detected. Shown is the percent of repeats with surviving cells of each strain in different cell densities, and the time to resume growth (color code). **(B)** *Msn4 expression increases along the growth curve in protein and transcript levels, while Msn2 expression remains stable:* expression was measured using fluorescent protein fusion (**B**, left), and transcription profiles (**B,** right). (**C).** *MSN4 expression is noisy while MSN2 expression follows the Poissonian variance:* mRNA molecules of *MSN2,4* were counted in > 4000 single cells with smFISH in exponentially growing cells (circles) and at OD_600_ = 4 (stars). Shown are the mean number of molecules at the x-axis and the Fano-factor (Left) and skewness (Middle) of the mRNA distribution at the y-axis. Dashed line represents the Poisson distribution parameters. (Right) smFISH imaging examples.

### Msn4 expression is environmental-sensitive and high noise

Msn4, the Msn2 duplicate, is also a stress genes activator^13,26^. Msn4-GFP was undetectable in reported measurements^34,35^, suggesting that its expression level is low during rapid growth. We reasoned that Msn4 expression increases along the growth curve to promote stress protection. This was indeed the case: Msn4 expression increased with cell density, both at the transcript and the protein levels (Figure 2B,C). This higher expression was accompanied by increased contribution to stress protection, as was measured by introducing H_2_O_2_ to strains deleted of msn2 in different cell densities (Figure 2A). Consistent with the control-noise tradeoff described above, this dynamic regulation of *MSN4* was accompanied by a high level of expression noise, which significantly exceeded the Poissonian variance (Figure 2C).

### Msn2 and Msn4 co-localize to the nucleus with the same dynamics in individual cells

Msn4 could collaborate with Msn2 in promoting stress protection by regulating the same set of genes or by inducing a distinct set of targets. Similarly, it could respond to the same, or to different sets of post-translational factors. Since activation of Msn2,4 culminates in nuclear localization^36,37^, we first followed the nuclear translocation dynamics of fluorescent-tagged Msn2 and Msn4 (Figure 3A). In response to stress, the two factors translocated to the nucleus within minutes, showing precisely the same kinetics within individual cells (Figure 3B,C-yellow shade, S2). Similarly, during the stochastic pulsing following stress^37,38^, translocation of the two factors was tightly synchronized within individual cells, but not between different cells (Figure 3C-pink shade). Deletion of one factor did not affect the dynamics of its duplicate (Figure S3).

**Figure. 3.**
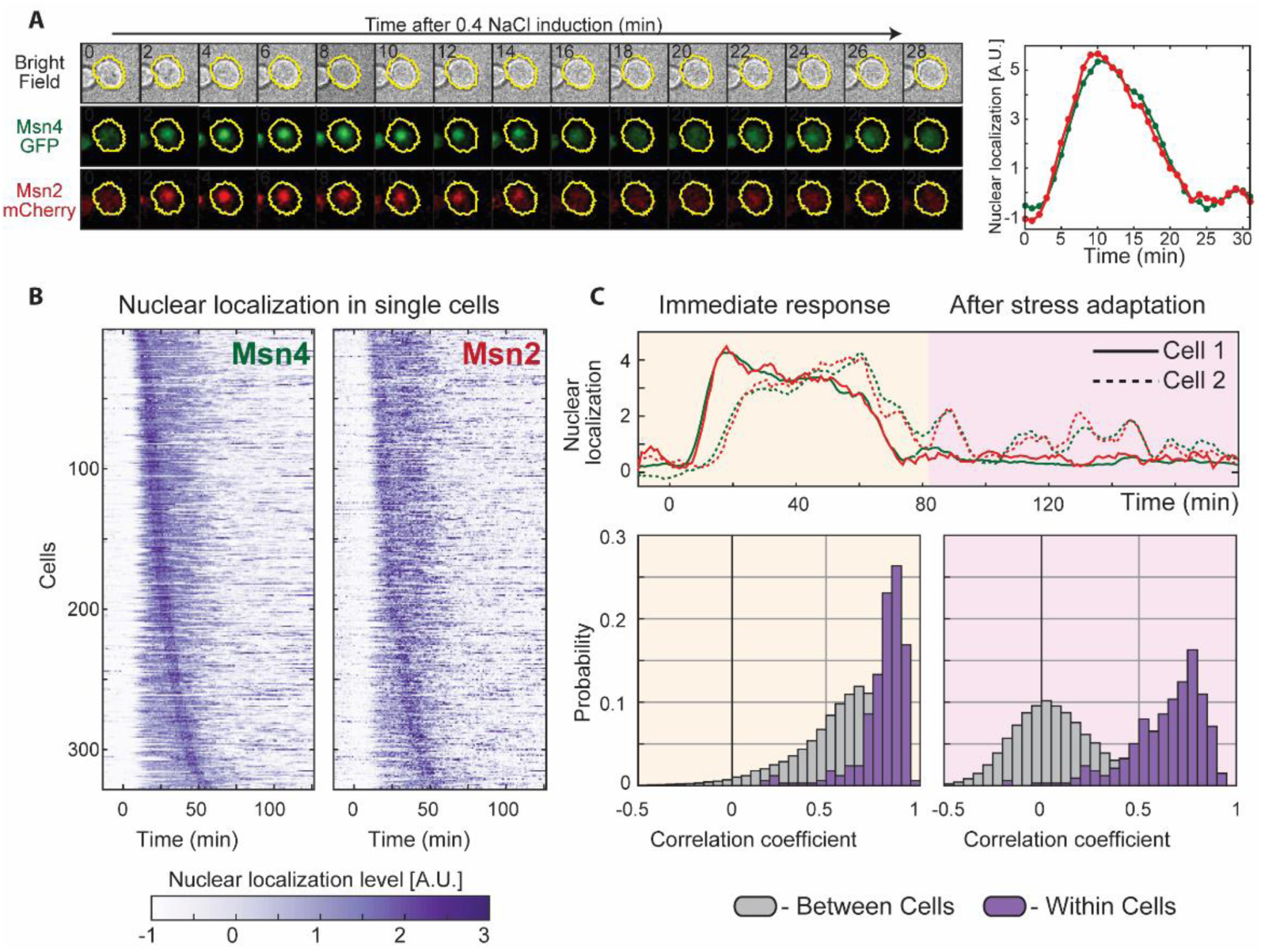
Msn2 and Msn4 translocate to the nucleus with the same dynamics in individual cells: Single cells expressing Msn4-GFP and Msn2-mChrerry were visualized using microfluidics-coupled live microscopy. Both proteins were readily visualized when cells were first cultured at intermediate or high OD (as MSN4 is undetectable in low ODs when cells grow exponentially). Cells were tracked as they were exposed to 0.4M, 1.2M or 1.4M NaCl. Cells were segmented and the nuclear localization of both proteins was quantified. **(A)** (left) A representative single cell in time in 3 channels. (right) quantification of the nuclear localization of Msn2 (red) and Msn4 (green). **(B)** Temporal traces of 328 single cells in 1.2M NaCl, ordered in both column by the time of Msn4-GFP nuclear localization. (0.4M/1.4M NaCl in Figure S2). **(C)** Correlations between the individual traces of Msn2 vs. Msn4 nuclear localization in single cells. Shown are the distributions of the correlation coefficients within the same (purple) or in different (gray) cells, separately comparing the immediate response (left) and the longer-time dynamics (right).

### Msn2 and Msn4 induce the same set of target genes

Next, we examined for differences in Msn2,4 target genes. In rapidly growing cells, deletion of *MSN2* strongly reduced stress gene induction, while deletion of *MSN4* had little, if any, effect (Figure 4A, S5). Swapping the *MSN2,4* promoters completely reversed the target induction capacity of these factors (Figure 4B, S6B). The identity of the targets remained the same: Msn4 driven by the *MSN2* promoter induced precisely the same targets normally induced by Msn2. When tested in conditions where both factors are expressed to equivalent amounts, the two factors induced the same set of genes (Figure 4C, S6). Since a previous study^39^ which followed stress gene induction using fluorescence reporters, indicated some differences in individual targets dependence on Msn2,4, we examined specifically the genes reported to be differently regulated. However, none of these genes showed any difference in their Msn2,4 dependency in any of the 6 conditions for which we performed tight time-course measurements (Figure S7). To further corroborate these results, we also measured the genome-wide binding profiles of the two factors, using the sensitive ChEC-seq method^40^. The two factors bound to the precise same promoters, occupied the precise same positions within individual promoters, and showed an identical preference for their common (known) DNA binding motif (Figure S8). This identity of Msn2,4 targets is consistent with the high conservation of their DNA binding domains (Figure 3D), and identity of their *in-vitro* DNA binding preferences^41^ (Figure S9). We conclude that Msn2,4 proteins are co-regulated by the same signals and at the same kinetics, and activate the same set of target genes with the same kinetics, essentially functioning as one TF.

**Fig. 4.**
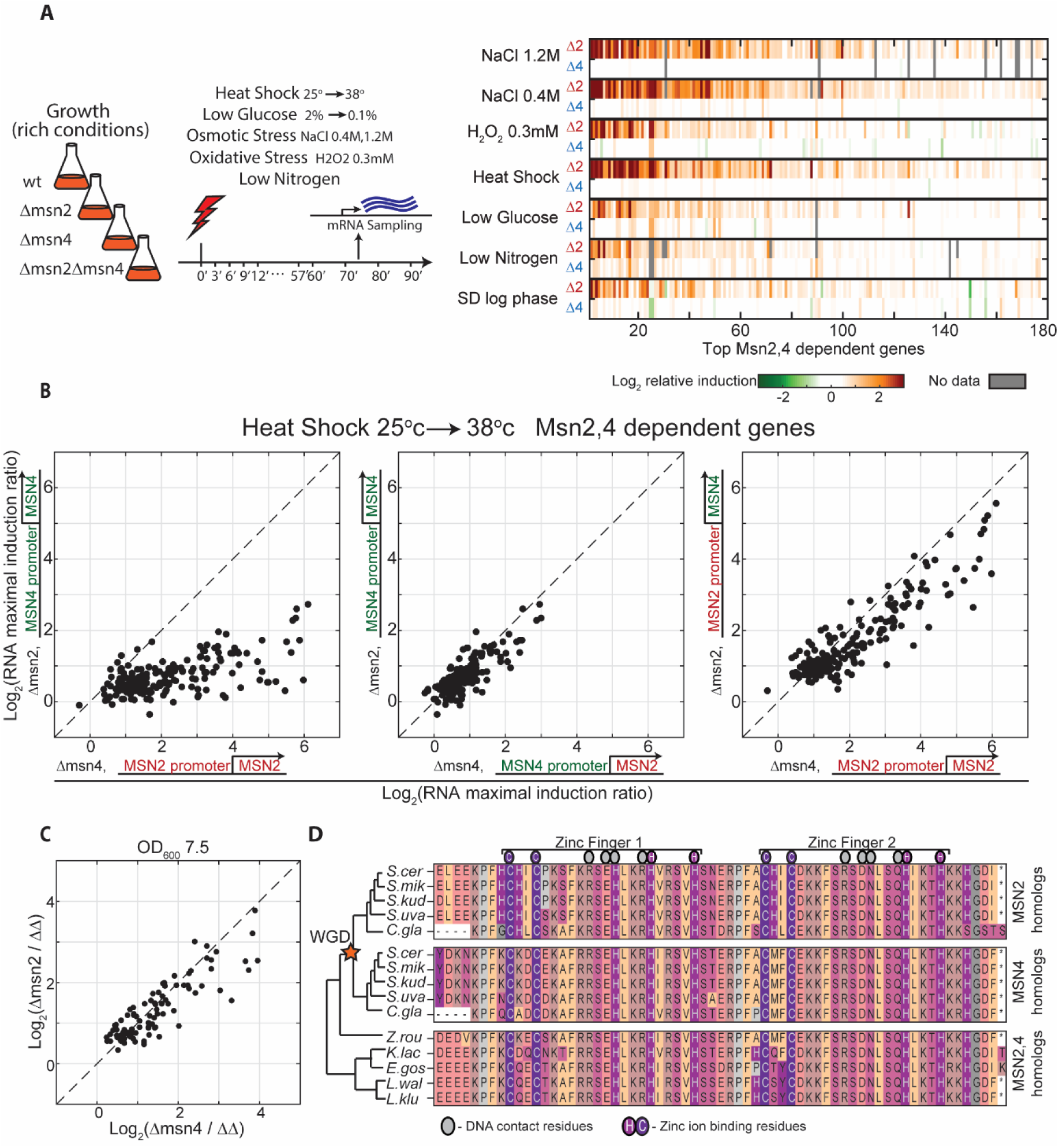
Redundancy in Msn2 and Msn4 activity: **(A)** *Stress response in rapidly growing cells depends on Msn2, but not Msn4:* Exponentially growing cells were exposed to the indicated stresses. (Left) Genome-wide transcription profiles were measured at 3’ time resolution following stress induction, for the first 60 minutes, and 10’ for the next 30 minutes. (Right) The stress response of each gene was summarized by its integrated (log2) change over the time course. The experiment was repeated in wild-type cells, in single deleted cells (Δmsn2, Δmsn4) and double deleted cells (Δmsn2Δmsn4). Shown are the differences between gene induction of the wild-type vs. the single deletion strains (Δ*msn2* or Δmsn4 at the top or bottom rows, respectively). 180 genes are shown, selected and ordered by the fold-change difference in their response in the wild-type vs. single deletion cells, these genes contain stress induced modules defined by other studies (Figure S4). **(B)** *Msn2 and Msn4 induce the same set of target genes:* during exponential growth, when Msn2 expression is higher than Msn4, deletion of Msn2 results in a significantly stronger effect on stress gene expression (left), but this effect was fully reversed by swapping the Msn2 and Msn4 promoters (middle and right). Each dot represents a target gene and its induction ratio between the indicated strain and the double msn2 and msn4 deletion strain. **(C)** In high OD (7.5), when both factors are expressed, stress genes are induced equally. Each dot is an induced target gene. (D) *Msn2 and Msn4 bind DNA through a highly conserved DNA binding domain:* Alignment of Msn2 and Msn4 DNA binding domains and their homologues in 10 species of the *Ascomycota* phylum that diverged before or after WGD (star). Colors indicate amino acid residue types.

### Differential design of the Msn2,4 promoters explains the differences in their expression flexibility and noise

Msn2 expression is stable along the growth curve, while Msn4 is strongly induced. To examine whether this differential regulation in expression is specific to these conditions, or is rather a more general property of the two genes, we surveyed a dataset composed of thousands of transcription profiles^12,42,43^. Expression of Msn2 showed little variability under all conditions tested, while Msn4 was highly variable (Figure 5A,B). Expression of Msn2 and Msn4 therefore conforms to the general tradeoff between expression noise and regulatory control: Msn2 is stable across conditions and shows low cell to cell variability (noise), while Msn4 expression readily responds to environmental signals and is noisy.

**Figure. 5.**
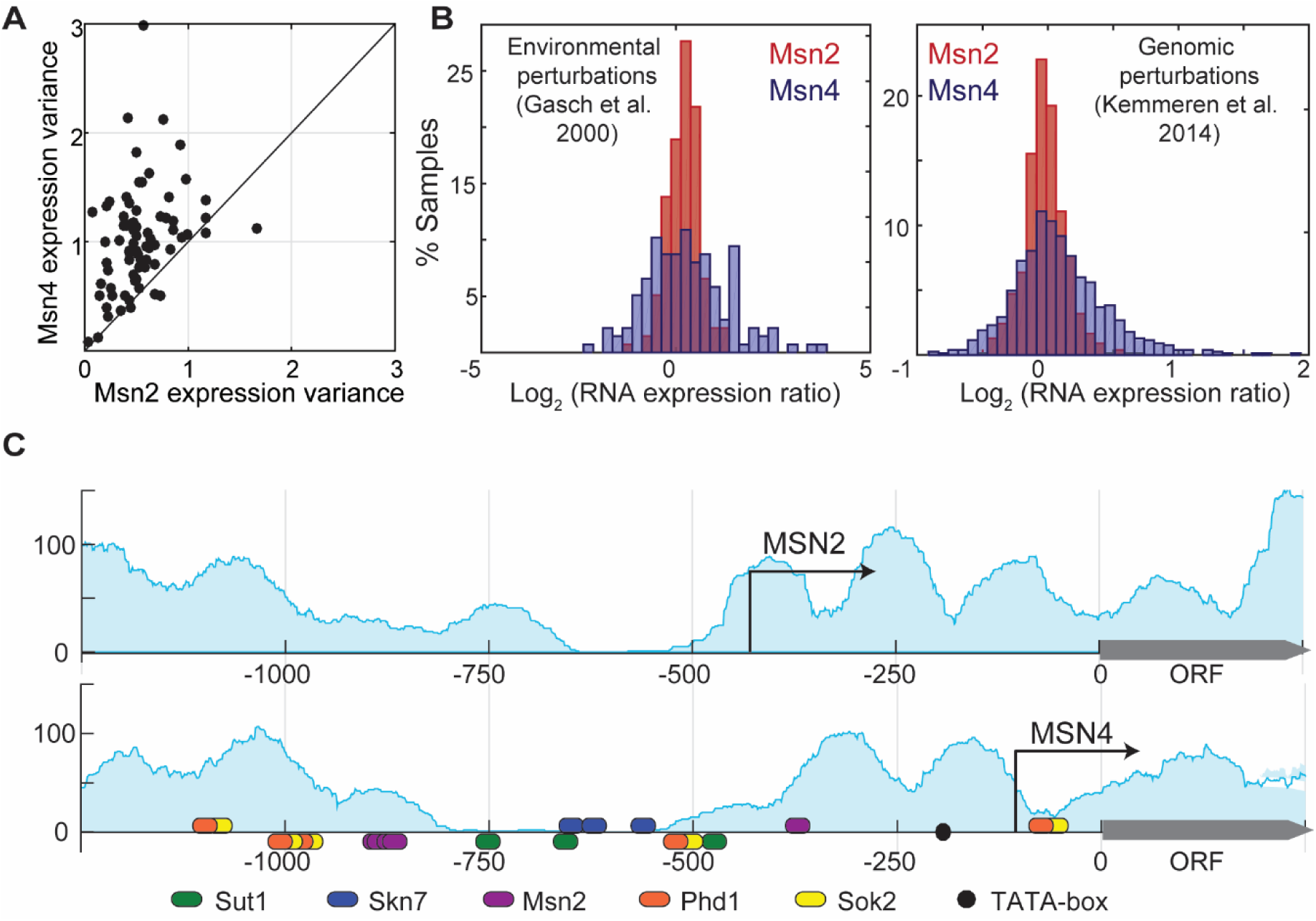
Msn2 expression is stable across conditions while Msn4 expression is variable. These properties are encoded in their promoters. (**A-B**) Three data types were considered. First, we downloaded >230 mRNA expression datasets available in SPELL^42^, and compared the variance of *MSN2* and *MSN4* expression in each dataset with more than 20 samples (A, each dataset is a dot). Second, we compared the distribution of *MSN2* and *MSN4* expression levels in two large datasets, representing multiple stress conditions^12^ (B, left) or gene deletions^43^(B, right). (**C**) *The MSN2 promoter displays properties of the stable, low noise type, while MSN4 promoter conforms to the flexible noisy type:* The pattern of nucleosome occupancy along the two promoters as defined by Weiner et al.^47^, is shown in blue shade. Arrows represent TSS positions, as defined by Park et al.^48^. Ellipses denote TFs binding sites as defined by MacIsaac et al.^49^ TATA-box (black circles) is defined as: TATA[AT]A[AT].

Previous studies have defined promoter designs that encode for flexible and noisy, or stable and low-noise expression^44–46^. Flexible promoters tend to contain a TATA box and bind nucleosomes immediately upstream to their Transcription Start Site (TSS), while stable promoters lack a TATA box and display a Nucleosome Free Region (NFR) upstream of their TSS. Consistent with their differential flexibility, we find that the *MSN4* promoter contains a TATA box, binds nucleosomes around its TSS and contains a large number of TF binding sites. By contrast, the *MSN2* promoter does not contain a TATA box, displays an NFR immediately upstream of the TSS and is largely devoid of TF binding sites (Figure 5C; data from^47–49^).

When aligned by their coding frames, the nucleosome patterns along the upstream regions of *MSN2* and *MSN4* promoters are highly similar. However, the location of the TSS is different: in *MSN4*, the TSS is positioned ∼105 bp away in a region that is nucleosome occupied, while in *MSN2* the TSS is significantly further upstream and located on the border of an NFR. The resulting 5’UTR of *MSN2* is exceptionally long (∼430 bp of length, found in only 2% of *S. cerevisiae* genes). Deleting this NFR region from the *MSN4* promoter, practically eliminated Msn4 induction along the growth curve. By contrast, deleting regions close to the ORF had little, if any further effect (Figure S10). Furthermore, replacing this region in the *MSN4* promoter by the corresponding region from *MSN2* promoter, including its new TSS and NFR, increased *MSN4* expression and reduced its noise (Figure S11). Therefore, this promoter region accounts for the differential expression characteristics of *MSN2* and *MSN4*.

### A shift in the TSS following WGD event modified MSN2 promoter design

Msn2,4 were duplicated in the whole-genome duplication (WGD) event, ∼100 million years ago^11^, and were retained in all WGD species tracing to this event. To examine whether the differential promoter structure of *MSN2,4* is conserved in other WGD species, we used available 5’ RNA data^50^ and further profiled TSS positioning in these species. The TSS positions of the *MSN2* and *MSN4* homologues were conserved in all post-WGD species (Figure 6A). Sequence analysis further indicated that the TATA box was conserved in all *MSN4* homologues but absent from all *MSN2* homologues (Figure 6A). We next profiled 5’ RNA in two non-WGD species. The transcript of the single *MSN* homologue has a short 5’UTR, similar to that of *MSN4*. This pattern of conservation is consistent with a scenario in which the stable *MSN2* promoter evolved from an ancestral flexible promoter through a shift in the TSS to a distant, TATA-lacking position, at the boundary of a nearby NFR.

**Figure. 6.**
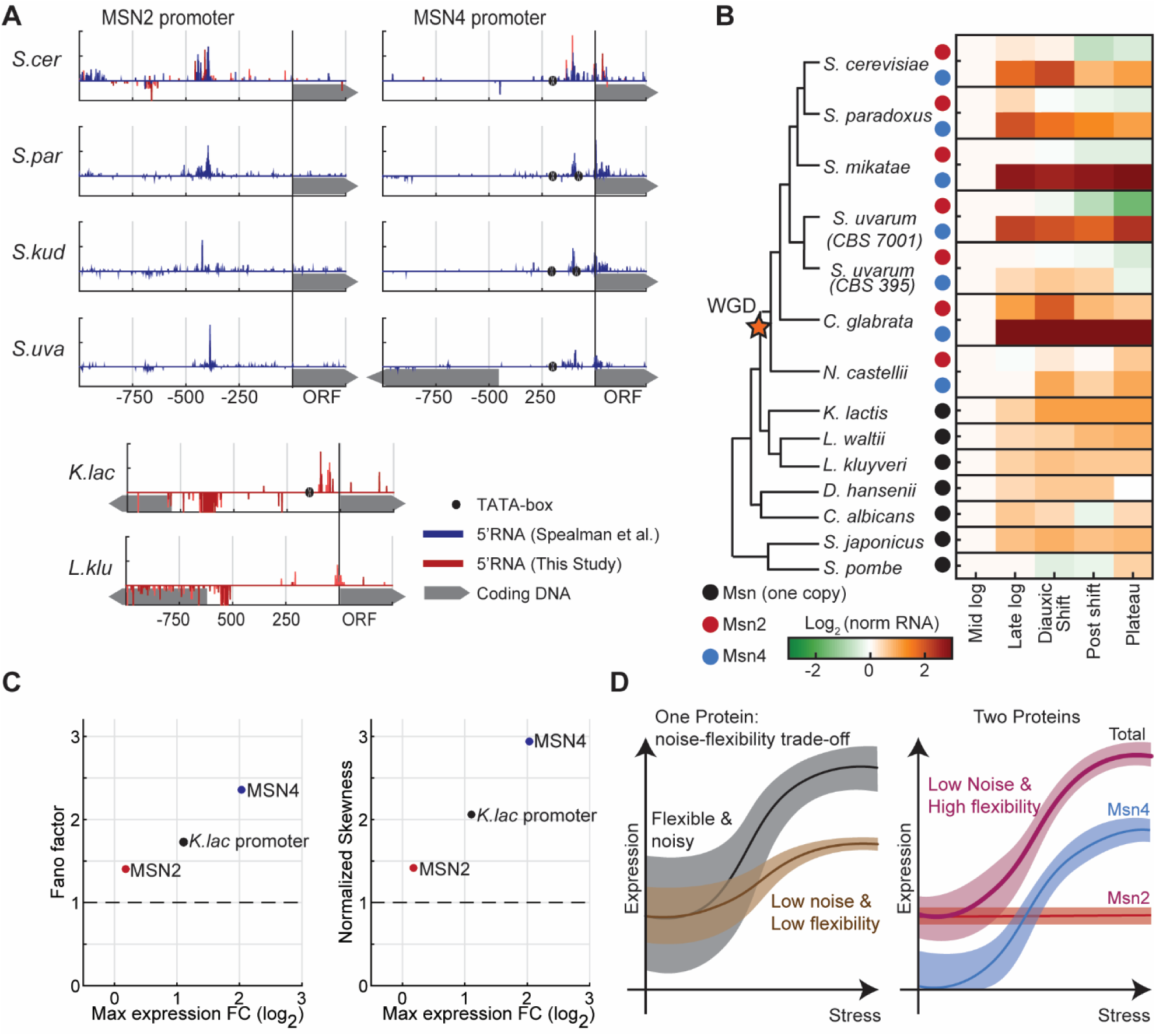
Msn2 shifted its TSS and gained a stable expression pattern in species that diverged from *S. cerevisiae* following WGD. **(A)** *The MSN2 promoter displays an uncharacteristically long 5’UTR that is conserved in all species that diverged after the WGD event:* Shown are the promoter maps of *MSN2,4* homologues in the indicated species. mRNA 5’ end mapping data from Spealman et al.^50^ is shown in blue, mRNA 5’ end from this study in red. TATA-box is defined as in figure 5C. **(B)** *MSN2 homologues are stably expressed along the growth curve, while MSN4 homologues show the flexible expression of the single MSN* homologues *found in species that diverged from S. cerevisiae prior to the WGD event:* Shown are expression levels of the Msn2,4 homologues in all indicated species, in 5 time-points along the growth curve. Data from Thompson et al.^51^ **(C)** *Expression of the K.lactis Msn2,4 homologue shows intermediate flexibility and noise.* On the x axis the maximal fold change expression of Msn2, Msn4 and the *K.lactis* homologue (data from: Thompson et al.^51^). Y axes show attributes of the expression distribution measured by smFISH, in Msn2, Msn4 and Msn2 in *S.cerevisiae* driven by the promoter of the *K.lactis* homologue. Shown are the fano factor (left) and the skewness of the distribution normalized to the skewness of a Poisson distribution with the same mean as the data (right). **(D)** *Model*: duplication of *Msn2,4 resolved conflict between environmental responsiveness and noise:* Single genes whose expression is sensitive to environmental conditions but will suffer from high noise in non-stressed conditions, limiting the ability to precisely tune intermediate expression levels while maintaining environmental-responsive expression. Gene duplication can resolve this conflict. See text for details.

### MSN4 accentuated the environmental-responsive but noisy expression of the non-WGD homologue, while MSN2 gained a stable, low-noise expression

To examine whether the differential expression flexibility of *MSN2,4* is also conserved in the other post-WGD species, we used available expression data^51^ of 13 yeast species along their growth curves. In all post-WGD species, *MSN4* expression increased along the growth curve, while *MSN2* expression remained stable (Figure 6B). The same dataset also profiled non-WGD species, allowing us to also examine the expression of the *MSN* single homologue in these species. The single *MSN* homologue in the non-WGD species showed a moderate induction along the growth curve, with dynamic range that was larger than that of *MSN2*, but lower than that of *MSN4* (Figure 6B). To examine whether this intermediate regulation is also reflected in the expression noise, we introduced the *MSN* promoter from *K. lactis*, a non-WGD species into *S. cerevisiae*, upstream of the *MSN2* ORF, and measured expression noise using smFISH. As predicted, this promoter showed an intermediate noise level that was higher than *MSN4* but lower than *MSN2* (Figure 6C). In fact, when plotted on the noise-control curve, the three promoters all fell on the same line, consistent with same-proportion change in noise and dynamic range of regulated expression. Therefore, our analysis suggests that *MSN2* gained its stable, low-noise expression following the duplication event, likely by shifting its TSS, while *MSN4* accentuated the regulated expression of the ancestral factor, likely through the acquisition of new binding sites for transcription factors, increasing its dynamic range and expression noise.

## Discussion

Taken together, our study defined a novel role for the Msn2,4 duplication. We were initially surprised to find that these two duplicates regulate the same set of target genes, translocate to the nucleus with the precise same dynamics, and are expressed in an overlapping set of conditions. What limits their replacement, in at least some species, by a single factor of a more refined transcriptional control? Our data shows that Msn2,4 function as one unit whose expression is both environmentally-responsive and low-noise (Figure 6D), thereby resolving an inherent conflict that limits the tuning of individual gene expression. Msn2 provides the low-noise basal expression, whereas Msn4 is induced when additional amounts are needed.

What could be the evolutionary force promoting this new evolution? Since the MSN duplication traces to the WGD event, it is tempting to propose that its new expression characteristics were driven by the shift in metabolism: Rapidly growing non-WGD species respire, while WGD species ferment. Following this metabolic change, genes needed in respiring cells may shift from being constitutively expressed, to being Msn-dependent, as was indeed reported^52^. We propose that changes in the identity of Msn2,4-dependent genes accentuated its phenotypic effects on growth and drove selection for increased precision of Msn2 expression.

Gene duplications is a major source of evolutionary innovation^4,5^ that greatly contributes to the expansion of transcription networks^2,3^. A surprisingly large fraction of TF duplicates, however, retained a conserved DNA binding domain and bind to the same DNA motif (Figure S12), suggesting limited divergence in regulatory targets. These, and other duplicates of apparent redundant function^53–56^ do not comply with the accepted models of neo- or sub-functionalization explaining duplicate advantage. Our study suggests a third model whereby duplicates with redundant biochemical properties realize dynamic properties that are not possible, or difficult to achieve using a single factor. In the case of Msn2,4, duplication resolved a conflict between regulatory control and noise. In other cases, interactions between the factors may define a circuit with dynamic properties not implementable by a single gene^55,57,58^. Further studies will define the relative contribution of such circuit-forming mechanisms in explaining the retention of TFs or other duplicates.

## Supporting information

Supplemental Inforamation

## Acknowledgments

We thank members of our lab for fruitful discussions and comments on the MS. We thank Nir Friedman and his group for their help and fertile discussions, especially to Daphna Joseph-Strauss. We thank Yoav Breuer for his help with constructing and performing one of the smFISH experiment. We thank Alon Appleboim for his support and suggestions. This work was supported by the ISF, and the Minerva Center.

## Notes

#### Summary of Updates

Typo in the text

## References

1. Charoensawan, V., Wilson, D. & Teichmann, S. A. Genomic repertoires of DNA-binding transcription factors across the tree of life. Nucleic Acids Res. 38, 7364–7377, (2010).

2. Weirauch, M. T. & Hughes, T. R. A Catalogue of Eukaryotic Transcription Factor Types, Their Evolutionary Origin, and Species Distribution. in Sub-cellular biochemistry 52, 25–73 (2011).

3. Lambert, S. A. et al. The Human Transcription Factors. Cell 172, 650–665 (2018).

4. Conant, G. C. & Wolfe, K. H. Turning a hobby into a job: How duplicated genes find new functions. Nat. Rev. Genet. 9, 938–950 (2008).

5. Soskine, M. & Tawfik, D. S. Mutational effects and the evolution of new protein functions. Nat. Rev. Genet. 11, 572–582 (2010).

6. Hittinger, C. T. & Carroll, S. B. Gene duplication and the adaptive evolution of a classic genetic switch. Nature 449, 677–681 (2007).

7. Des Marais, D. L. & Rausher, M. D. Escape from adaptive conflict after duplication in an anthocyanin pathway gene. Nature 454, 762–765 (2008).

8. Voordeckers, K., Pougach, K. & Verstrepen, K. J. How do regulatory networks evolve and expand throughout evolution? Curr. Opin. Biotechnol. 34, 180–188 (2015).

9. Pérez, J. C. et al. How duplicated transcription regulators can diversify to govern the expression of nonoverlapping sets of genes. Genes Dev. 28, 1272–7 (2014).

10. Baker, C. R., Hanson-Smith, V. & Johnson, A. D. Following gene duplication, paralog interference constrains transcriptional circuit evolution. Science 342, 104–8 (2013).

11. Wolfe, K. H. & Shields, D. C. Molecular evidence for an ancient duplication of the entire yeast genome. Nature 387, 708–713 (1997).

12. Gasch, A. P. et al. Genomic Expression Programs in the Response of Yeast Cells to Environmental Changes. Mol. Biol. Cell 11, 4241–4257 (2000).

13. Schmitt, A. P. & McEntee, K. Msn2p, a zinc finger DNA-binding protein, is the transcriptional activator of the multistress response in Saccharomyces cerevisiae. Proc. Natl. Acad. Sci. U. S. A. 93, 5777–82 (1996).

14. Estruch, F. Stress-controlled transcription factors, stress-induced genes and stress tolerance in budding yeast. FEMS Microbiol. Rev. 24, 469–486 (2000).

15. Elowitz, M. B., Levine, A. J., Siggia, E. D. & Swain, P. S. Stochastic gene expression in a single cell. Science 297, 1183–6 (2002).

16. Raser, J. M. & O’Shea, E. K. Noise in gene expression: origins, consequences, and control. Science 309, 2010–3 (2005).

17. Lehner, B. Selection to minimise noise in living systems and its implications for the evolution of gene expression. Mol. Syst. Biol. 4, (2008).

18. Metzger, B. P. H., Yuan, D. C., Gruber, J. D., Duveau, F. & Wittkopp, P. J. Selection on noise constrains variation in a eukaryotic promoter. Nature 521, 344–347 (2015).

19. Eldar, A. & Elowitz, M. B. Functional roles for noise in genetic circuits. Nature 467, 167–173 (2010).

20. Raj, A. & van Oudenaarden, A. Nature, Nurture, or Chance: Stochastic Gene Expression and Its Consequences. Cell 135, 216–226 (2008).

21. Yaakov, G., Lerner, D., Bentele, K., Steinberger, J. & Barkai, N. Coupling phenotypic persistence to DNA damage increases genetic diversity in severe stress. Nat. Ecol. Evol. 1, 0016 (2017).

22. Newman, J. R. S. et al. Single-cell proteomic analysis of S. cerevisiae reveals the architecture of biological noise. Nature 441, 840–846 (2006).

23. Lehner, B. Conflict between Noise and Plasticity in Yeast. PLoS Genet. 6, e1001185 (2010).

24. Hornung, G. et al. Noise-mean relationship in mutated promoters. Genome Res. 22, 2409–17 (2012).

25. Choi, J. K. & Kim, Y.-J. Intrinsic variability of gene expression encoded in nucleosome positioning sequences. Nat. Genet. 41, 498–503 (2009).

26. Martínez-Pastor, M. T. et al. The Saccharomyces cerevisiae zinc finger proteins Msn2p and Msn4p are required for transcriptional induction through the stress response element (STRE). EMBO J. 15, 2227–2235 (1996).

27. Raj, A., van den Bogaard, P., Rifkin, S. A., van Oudenaarden, A. & Tyagi, S. Imaging individual mRNA molecules using multiple singly labeled probes. Nat. Methods 5, 877–879 (2008).

28. Munsky, B., Neuert, G. & van Oudenaarden, A. Using gene expression noise to understand gene regulation. Science 336, 183–7 (2012).

29. Bar-Even, A. et al. Noise in protein expression scales with natural protein abundance. Nat. Genet. 38, 636–643 (2006).

30. Fraser, H. B., Hirsh, A. E., Giaever, G., Kumm, J. & Eisen, M. B. Noise Minimization in Eukaryotic Gene Expression. PLoS Biol. 2, e137 (2004).

31. Wang, Z. & Zhang, J. Impact of gene expression noise on organismal fitness and the efficacy of natural selection. Proc. Natl. Acad. Sci. U. S. A. 108, E67–76 (2011).

32. Richard, M. & Yvert, G. How does evolution tune biological noise? Front. Genet.5, 374 (2014).

33. Keren, L. et al. Massively Parallel Interrogation of the Effects of Gene Expression Levels on Fitness. Cell 166, 1282–1294.e18 (2016).

34. Huh, W.-K. et al. Global analysis of protein localization in budding yeast. Nature 425, 686–691 (2003).

35. Breker, M., Gymrek, M., Moldavski, O. & Schuldiner, M. LoQAtE—Localization and Quantitation ATlas of the yeast proteomE. A new tool for multiparametric dissection of single-protein behavior in response to biological perturbations in yeast. Nucleic Acids Res. 42, D726–D730 (2014).

36. Petrenko, N., Chereji, R. V., McClean, M. N., Morozov, A. V. & Broach, J. R. Noise and interlocking signaling pathways promote distinct transcription factor dynamics in response to different stresses. Mol. Biol. Cell 24, 2045–2057 (2013).

37. Hao, N., Budnik, B. A., Gunawardena, J. & O’Shea, E. K. Tunable signal processing through modular control of transcription factor translocation. Science 339, 460–4 (2013).

38. Lin, Y., Sohn, C. H., Dalal, C. K., Cai, L. & Elowitz, M. B. Combinatorial gene regulation by modulation of relative pulse timing. Nature 527, 54–58 (2015).

39. AkhavanAghdam, Z., Sinha, J., Tabbaa, O. P. & Hao, N. Dynamic control of gene regulatory logic by seemingly redundant transcription factors. Elife 5, (2016).

40. Zentner, G. E., Kasinathan, S., Xin, B., Rohs, R. & Henikoff, S. ChEC-seq kinetics discriminates transcription factor binding sites by DNA sequence and shape in vivo. Nat. Commun. 6, 8733 (2015).

41. Siggers, T., Reddy, J., Barron, B. & Bulyk, M. L. Diversification of Transcription Factor Paralogs via Noncanonical Modularity in C2H2 Zinc Finger DNA Binding. Mol. Cell 55, 640–648 (2014).

42. Hibbs, M. A. et al. Exploring the functional landscape of gene expression: directed search of large microarray compendia. Bioinformatics 23, 2692–2699 (2007).

43. Kemmeren, P. et al. Large-Scale Genetic Perturbations Reveal Regulatory Networks and an Abundance of Gene-Specific Repressors. Cell 157, 740–752 (2014).

44. Field, Y. et al. Distinct Modes of Regulation by Chromatin Encoded through Nucleosome Positioning Signals. PLoS Comput. Biol. 4, e1000216 (2008).

45. Tirosh, I. & Barkai, N. Two strategies for gene regulation by promoter nucleosomes. Genome Res. 18, 1084–1091 (2008).

46. Nicolas, D., Phillips, N. E. & Naef, F. What shapes eukaryotic transcriptional bursting? Mol. Biosyst. 13, 1280–1290 (2017).

47. Weiner, A. et al. High-Resolution Chromatin Dynamics during a Yeast Stress Response. Mol. Cell 58, 371–386 (2015).

48. Park, D., Morris, A. R., Battenhouse, A. & Iyer, V. R. Simultaneous mapping of transcript ends at single-nucleotide resolution and identification of widespread promoter-associated non-coding RNA governed by TATA elements. Nucleic Acids Res. 42, 3736–3749 (2014).

49. MacIsaac, K. D. et al. An improved map of conserved regulatory sites for Saccharomyces cerevisiae. BMC Bioinformatics 7, 113 (2006).

50. Spealman, P. et al. Conserved non-AUG uORFs revealed by a novel regression analysis of ribosome profiling data. Genome Res. 28, 214–222 (2018).

51. Thompson, D. A. et al. Evolutionary principles of modular gene regulation in yeasts. 2, 603 (2013).

52. Tirosh, I., Wong, K. H., Barkai, N. & Struhl, K. Extensive divergence of yeast stress responses through transitions between induced and constitutive activation. Proc. Natl. Acad. Sci. U. S. A. 108, 16693–8 (2011).

53. DeLuna, A. et al. Exposing the fitness contribution of duplicated genes. Nat. Genet. 40, 676–681 (2008).

54. Dean, E. J., Davis, J. C., Davis, R. W. & Petrov, D. A. Pervasive and Persistent Redundancy among Duplicated Genes in Yeast. PLoS Genet. 4, e1000113 (2008).

55. Kafri, R., Springer, M. & Pilpel, Y. Genetic Redundancy: New Tricks for Old Genes. Cell 136, 389–392 (2009).

56. Diss, G. et al. Gene duplication can impart fragility, not robustness, in the yeast protein interaction network. Science 355, 630–634 (2017).

57. Alon, U. Network motifs: theory and experimental approaches. Nat. Rev. Genet. 8, 450–461 (2007).

58. Teichmann, S. A. & Babu, M. M. Gene regulatory network growth by duplication. Nat. Genet. 36, 492–496 (2004).

59. Rahman, S. & Zenklusen, D. Single-molecule resolution fluorescent in situ hybridization (smFISH) in the yeast S. cerevisiae. Methods Mol. Biol. 1042, 33–46 (2013).

60. Avraham, N., Soifer, I., Carmi, M. & Barkai, N. Increasing population growth by asymmetric segregation of a limiting resource during cell division. Mol. Syst. Biol. 9, (2013).

61. Kafri, M., Metzl-Raz, E., Jona, G. & Barkai, N. The Cost of Protein Production. Cell Rep. 14, 22–31 (2016).

62. Orsi, G. A., Kasinathan, S., Zentner, G. E., Henikoff, S. & Ahmad, K. Mapping Regulatory Factors by Immunoprecipitation from Native Chromatin. in Current Protocols in Molecular Biology 110, 21.31.1-21.31.25 (John Wiley & Sons, Inc., 2015).

63. Pelechano, V., Wei, W. & Steinmetz, L. M. Extensive transcriptional heterogeneity revealed by isoform profiling. Nature 497, 127–131 (2013).

64. Ihmels, J., Bergmann, S. & Barkai, N. Defining transcription modules using large-scale gene expression data. Bioinformatics 20, 1993–2003 (2004).

65. De Boer, C. G. & Hughes, T. R. YeTFaSCo: A database of evaluated yeast transcription factor sequence specificities. Nucleic Acids Res. 40, D169–D179 (2012).

66. Yofe, I. et al. One library to make them all: streamlining the creation of yeast libraries via a SWAp-Tag strategy. Nat. Methods 13, 371–378 (2016).

