## Supplemental Inforamation for "Resolving noise-control conflict by gene duplication"

**This file includes:**

Materials and Methods
Figs. S1 to S12
Tables S1 to S3
References for supplementary Materials

### Materials and Methods

Strains

All strains used in this study and their genotypes are listed in Table S1. All the strains were constructed by standard genetic methods and were validated by PCR and/or sequencing of the relevant DNA.

Single molecule Fluorescence In-Situ Hybridization

For each gene (MSN2 and MSN4), a set of 48 probes was generated as described in Raj et al.[27] The probes were designed by the online program Stellaris™ Probe Designer from Biosearch Technologies, and were ordered with a fluorescent dye CAL Fluor Red® 590. Probe sequences are listed in Tables S2-S3.

Cells were grown overnight in synthetic complete (SC) medium at 30° C and constant shaking. Then diluted to reach the wanted cell densities after ~12 hours. Cells were fixated, prepared and hybridized as described in Rahman et al. [59]

Images were acquired with 100x 1.4 oil UPLSAPO objective, using Olympus IX83 based Live-Imaging system equipped with CSU-W1 spinning disc (sCMOS digital Scientific Grade Camera 4.2 MPixel). For each sample 4-6 different positions were chosen. In each position three Z-stacks images were taken with step size of 200nm for a total of >6µm: Bright-field image- 488nm laser with 100mW, DAPI image- 405nm laser with 120mW and exposure time of 250ms , mRNA image- 561nm laser with 100mW and exposure time of 1000ms. Each z-plane image was of size 2048x2048 pixels.

*Single molecule quantification*- Cells were segmented using a modification of a custom MATLAB software[60]. In this modification, cell centers were defined manually using the bright field images and cells borders were found automatically. mRNA counts were then performed for each cell based on the custom made MATLAB software from Raj et al.[27]

Stress experiments for RNA-seq levels

In these experiments we used the wt strains, the single msn2 or msn4 deletion strains, and the double msn2,msn4 deletion strain. Some of the experiments were also done with the strains with swapped promoters: MSN2 ORF under MSN4 promoter with a deletion of msn4, and the opposite- MSN4 ORF under MSN2 promoter, with an msn2 deletion).

*Growth Conditions* - Cells were grown overnight in rich medium – YPD or synthetic complete (SC) medium at 30° C (unless otherwise noted), then cells were diluted and exponentially grew for 6-8 hours before introducing the stress:

*Oxidative Stress* - Cells were grown continuously in 30° C. $H_2O_2$ was added to a final concentration of 0.3mM.

*Heat Shock* - Cells were grown continuously in 25° C, then cell culture was moved to a new flask located inside a bath orbital shaker (Cat. WBT-450, MRC) preheated to 37° C. It took less than 90 seconds for the culture to reach 37° C.

*Glucose limitation* - Cells were grown in SC medium with 2% glucose (Sigma-Aldrich). Then cells were washed twice and resuspended in SC with 0.1% glucose. Samples were taken before the washes (2% glucose), after every wash and for the next 100 min.

*Osmotic Shock* - Cells were grown continuously in 30° C. 4M NaCl solution was added to the culture to a final concentration of 0.4 M or 1.2M.

*Low nitrogen* - Cells were grown in SC medium with 2% glucose (Sigma-Aldrich). Then cells were washed twice and resuspended in Nitrogen-depleted medium (0.67% Yeast Nitrogen Base without amino acids and ammonium sulfate (Bacto-YNB), 2% glucose, 0.05 mM ammonium sulfate, 20 mg/l Urcail, 20 mg/l Histidine, 100 mg/l Leucin, 20 mg/l Methionine).

*Growth into stationary phase* – Cells were grown in SC in 30° C without changing the media.

RNA sample collection, extraction and sequencing levels

Cells were grown overnight to stationary phase and then diluted in 100ml to reach OD600 of 0.2-0.4 after 6-8 hours in constant shaking. A sample for time point zero reference was taken, then we introduced a stress perturbation as described above. For growing into stationary phase experiment, a sample of 1ml was taken every 20/30 minutes. For all other conditions a sample of 1.5ml was collected every 3 minutes for the first hour, and every 10 minutes for additional half/one hour. Samples were immediately centrifuged for 40 seconds in 13,000 rpm. The supernatant was removed, pellets were frozen in liquid nitrogen and stored at −80° C until RNA preparation.

RNA was extracted using modified protocol of nucleospin® 96 RNA kit. Specifically, cell lysis was done in a 96 deep-well plate by adding 450μl of lysis buffer containing 1M sorbitol (Sigma-Aldrich), 100mM EDTA and 0.45μl lyticase (10IU/μl). The plate was incubated at 30° C for 30 minutes in order to break the cell wall, and then centrifuged for 10 minutes at 2500 rpm, and supernatant was removed. From this stage, extraction proceeded as in the protocol of nucleospin® 96 RNA kit, only substituting β-mercaptoethanol with DTT.

For all samples sequenced by Illumina HiSeq 2500, RNA libraries were created as follows: Fragmented, poly(A)-selected RNA extracts of ∼200 bp size were reverse-transcribed to cDNA using barcoded poly(T) primers. cDNA was amplified and sequenced with Illumina HiSeq 2500 using a primer complementary to the opposite adaptor to the poly(A).

For all samples sequenced by Illumina NextSeq 500, RNA libraries were created as follows: poly(A) RNA was selected by reverse transcription with a barcoded poly(T) primer. The barcoded DNA-RNA hybrids were pooled and fragmented by a hyperactive variant of the Tn5 transposase. Tn5 was stripped off the DNA by treatment with SDS 0.2% followed by SPRI cleanup and the cDNA was amplified and sequenced with Illumina NextSeq 500.

Processing and analysis of RNA-seq data

We mapped 50bp reads of the RNA-seq of every sample to the *S.cereviciae* genome (R64 in SGD) using bowtie (parameters: –best -a -m 2 -strata -5 10). After alignment to the genome, samples that had less than 150,000 reads were discarded from the analysis in order to prevent an artificial enrichment for highly expressed genes. The expression at those time points was calculated as the mean between the two closest time points in the time course. For every sequence we normalized for PCR bias using the unique molecular identifier (UMI), scoring each position on the genome by the unique number of UMI's it had out of all possible UMI's. For each gene we summed all the reads aligned to 400bp upstream its 3' end to 200bp downstream, in order to get the total expression of that gene. Reads that were aligned non-uniquely were split between the aligned loci according to the ratio of all other uniquely mapped reads in these regions. Number of reads for each sample was normalized to $10^6$.

Msn2-Promoter library preparation

We used 140 synthetic promoters from Keren et al.[33] , pooled together and transformed them to replace the native *MSN2* promoter, in a strain with Msn2 tagged with YFP and deleted of msn4. We collected ~200 colonies after the transformation, and measured YFP fluorescence with flow cytometer (BD LSRII system from BD Biosciences). We picked 50 strains that spanned the expression range and was highly similar between the different repeats.

Growth experiment in harsh stress

*MSN2 promoter library strains* – we grew the cells to stationary phase in SC media in 96-well plate under constant shaking and 30°c. Next, we diluted the cells with fresh SC media, in deep well plates with one glass bead in each well to generate proper shaking, so they will reach the wanted OD in the next morning. Then, right before stressing the cells, we took 150µl to measure ODs (using infinite200 reader, Tecan Inc.), and 100µl to the a flow cytometer to measure Msn2-YFP fluorescence.

*Single and double deletion trains* – we grew cells overnight to stationary phase in SC media and constant shaking, at 30°c. Next, we serial dilute the cells with fresh SC media in a 96-well plate to reach sequential different ODs in the next morning. Then, right before stressing the cells, we took 150µl to measure ODs (using infinite200 reader, Tecan Inc.)

*Stress and growth measurements* – we took 30µl of growing cells into plates with 120µl of H2O2. We inserted the plates into an automated handling robot (EVOware, Tecan Inc.) where cells were grown under constant shaking and 30°c. The robot was programed to take the plates out of the incubator every 45 minutes, vortex the plates, and measure the OD (using infinite200 reader, Tecan Inc.). Experiment lasted for ~70 hours.

*Growth analysis*- data from the growth measurements was parsed and processed. Time to exponential growth was calculated as the time of the maximal slope of the OD measurements in time. We calculated the median time and the std of the repeats.

### Competition experiment in SC media

Cells were grown O.N to stationery phase in SC. Then diluted and grown ~8 hours in exponential growth. Each strain of the Msn2-promoter library strains were then co-incubated with wt-mCherry strain at 30˚C. Wt initial frequency was ~50%. Every ~8 hours cells were diluted with fresh SC media so they will grow exponentially at all times. In addition, a sample was taken to measure OD, and to a flow cytometer to measure frequencies of each population. Flow cytometry measurements and analysis were done using BD LSRII system (BD Biosciences). Flow cytometer was conducted with excitation at 488nm and emission at 525±25nm for GFP samples. For mCherry markers, excitation was conducted at 594nm and emission at 610±10nm. The number of generations was calculated from the dilution factor. % of wt division rate was calculated as previously described in kafri et al.[61]

### Msn2,4 protein expression by flow cytometry

For this experiment, we used strain with Msn2 tagged with GFP, and a strain with Msn4 tagged with GFP and Msn2 tagged with mCherry. We grew the cells overnight in 5 mL SC media and 30°C to reach stationary phase, then to reach $OD_{600}$ ~0.4 after ~8 hours. Then, we serial diluted the cells in 96-well plate by diluting each column to the next one with 1:1 ratio with SC media, ending up with 120μl in each well, and 1:2 ratio of cells in adjacent columns. After overnight incubation in 30°C, and constant shaking, we measured fluorescence using flow cytometer.

Flow cytometry measurements and analysis were done using BD LSRII system (BD Biosciences). Flow cytometer was conducted with excitation at 488nm and emission at 525±25nm for GFP samples. For mCherry markers, excitation was conducted at 594nm and emission at 610±10nm. The average number of cells analyzed was 50,000. For the samples with high OD, 100μl of DDW was added to the sample before reading it in the FACS.

We calculated cell density using the flow cytometer parameters and output. This measure was calculated as following: $\frac{N}{V}C$ where R = flow rate (μl /sec), T = total flow time (sec), V = R*T = total volume read (μl), N = number of cells read (cells), C = dilution fix constant (values are either 1 for no dilution with DDW, or 1.8333 for samples that were diluted with 100 μl DDW).

We filtered G1 cells by the width size measure FSC-W, which has a bimodal distribution that corresponds to cells in G1 (smaller) and cells in G2/M (bigger). Next, we filtered outliers by two area measures – FSC-A and SSC-A. We used linear regression to describe FSC-A with SSC-A, and removed cells that were far from the regression line. We then applied linear regression to describe SSC-A with FSC-A and removed outliers in a similar manner.

In order to eliminate the background fluorescence we used a linear regression model that predicts background fluorescence. The independent variables were size parameters (FSC-W and SSC-W) and the cell density of the population. The dependent variable was the background fluorescence (GFP/mCherry). Model was trained on BY4741 cells with no fluorescent markers, then used to predict background in the other strains. Predicted background was subtracted from observed fluorescence for each cell.

### Time laps microscopy experiment

We used a strain with both Msn2-mCherry and GFP-Msn4 and a strain with GFP-Msn4 and a deletion of Msn2. We grew the cells overnight in SC media to reach stationary phase, then diluted them to reach the desired OD600 of ~7 after ~8 hours. When reaching the desired OD, cells were transferred to a microfluidics plate (catalog number: Y04C-02-5PK) for haploid yeast cells. We used ONIX CellAsic microfluidics system that allows changing the cells' media at a fast rate, in a predefined set time, while not interfering with the imaging process. During imaging, after ~15 minutes of normal media, a media with NaCl was added to the cells (0.4/1.2/1.4 M). Every minute, for each position on the plate, three images were taken – bright-field image, GFP image and mCherry image. Two positions were taken for each strain. Imaging under stress lasted for 4-8 hours.

We used a Zeiss AxioObserverZ1 inverted microscope equipped with Hamamatsu Flash4 sCMOS cameras. In every imaging instance, three images were taken: Bright-field image, GFP image using GFP filter with 20% intensity of HSP120 V lamp and with exposure time of 200 ms, and mCherry image using mPlum filter with same intensity and exposure as GFP. we used 2x2 binning resulting in 1024x1024 pixels of image size.

### Processing microscopy images and estimate nuclear localization levels

*Tracking and segmentation* - All images were subsequently analyzed using custom MATLAB software that segments and tracks individual cells along the movie in each bright field image frame, as previously described[60]. Briefly, cell borders were detected automatically in the last frame. Then the program goes back to the beginning of the experiment frame-by-frame, and for each cell in the image uses the centroid coordinates of the cells from the previous frame. Each centroid is expended until the borders of the cell in the current frame if found. The program also outputs a score for the segmentation that was used to filter out cells with low quality segmentation.

*Image processing* - Median filter: we ran 3x3 median filter on all GFP and mCherry images. Background removal was done by running mean filter of 50x50 on each image, then subtracting the filtered image from the original one. Rare events of missing frames (mainly due to focus issues) were interpolated to be the mean of the two adjacent frames.

*Calculating nuclear localization measure* - Our method uses image filtering with a filter shaped like a nucleus with radius of 3 pixels. We run the filter on each cell GFP/mCherry track, and then find the maximal coordinate of the filtered cell image, defining it as the center of the hypothetical nucleus. Our measure is the average over the pixels in the hypothetical nucleus divided by the pixels average in the hypothetical cytoplasm. For normalization, we divide each cell's localization in time by the minimal value for this cell. As a result, this method in fact gives signal-to-noise ratio. In order to align the GFP and the mCherry tracks together, we used z-score normalization (subtracted the mean and divided by standard deviation for each cell).

*Filtering bad cells*- we filtered out "bad" cells in two rounds, once after running segmentation, and once after calculating the dynamical attributes. In the first round, we filtered cells that answered one or more of the following conditions: (1) Cells with area outside the range defined $Median \pm 3 \cdot MAD$ (MAD is the mean absolute deviation) over all cells, at least 10% of the time. (2) Cells with segmentation score <8 at least 10% of the time. (3) Cells that did not appear from the beginning of the experiment. In the second round we removed cells with response amplitude below 1.1 or above 6.

### MSN2,4 homologues expression in growth

Data was taken from Thompson et al.[51] For each yeast species in this experiment, 5 time-points were measured as a ratio to mid-log sample: lag, late-log, diauxic-shift, post diauxic shift, plateau. For each time point 3 repeats were made. We show the average of the repeats. We exclude the first time-point (lag) from the figure due to a large number of missing data and repeats.

### 5'mRNA sequencing

Cells were grown overnight to stationary phase in SC media in 30°C, and then diluted and exponentially grew for ~12 hours in constant shaking. Samples were fixed by mixing with cold (-80°C) methanol. RNA was Poly(A)-selected, reverse transcribed to cDNA and barcoded at the 5' end using Dynabeads® Oligo(dT)$_{25}$ magnetic beads. cDNA was pooled and fragmented by a hyperactive variant of the Tn5 transposase. Tn5 was stripped off the DNA by SDS 0.2% treatment followed by SPRI cleanup and the cDNA was amplified and sequenced with Illumina NextSeq 500. Number of reads for each sample was normalized, and genomic tracks were created from the sequence reads, representing the enrichment on each position of the genome.

### CHeC-seq strains

We fused Msn2 or Msn4 to MNase (Amino Acids 83-231) using pGZ108 (pFA6a-3FLAG-MNase-kanMX6). This plasmid was a gift from Steven Henikoff (Addgene plasmid # 70231).

### CHeC-seq experiment

Cells were grown overnight to stationary phase in SC media in 30°C, and then diluted and grew for ~15 hours in 30°C and constant shaking until they reached OD600 of ~4. Then. CHeC-seq was performed as described in Zentner et al.[40] with 30 seconds of activated Mnase, and changes in the ethanolic precipitation (1h in -80°C), and SPRI beads size selection (0.8x). Library preparation was performed as describe in Orsi et al.[62] , with converting the S-300 column cleanup following the phenol-chloroform step to ethanolic precipitation. Libaries was sequenced with Illumina NextSeq 500.

CHeC-seq analysis

Reads were mapped to the S.cereviciae genome (R64 in SGD) using bowtie2. The first nucleotide of every read was counted as binding signal. All samples had >$10^6$ reads. Each sample was normalized to $10^7$ reads. Promoter length was defined as 700 base pairs upstream to the transcription start site or the distance to the upstream transcript (the shorter between these two). Transcription start and end sites were taken from Pelechano et al.[63] For the motif analysis, the average of the sum of signal of each 7-mer appearance (±10bp) in all of the promoter regions was calculated.

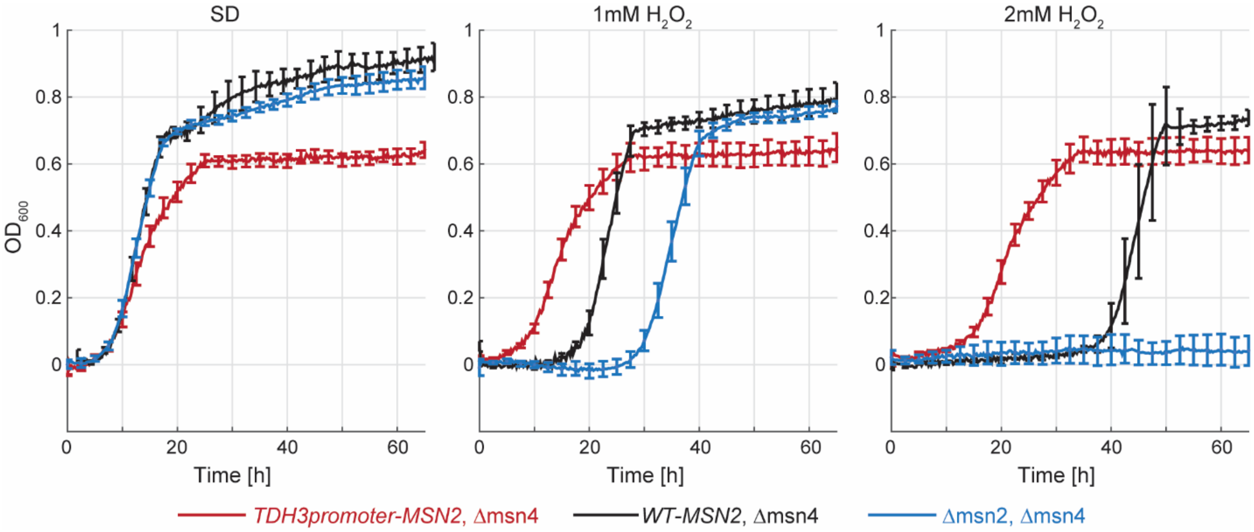


**Fig. S1.** *Growth curves in SD or $H_2O_2$ in Msn2 over-expression or deletion strains.* Cells were grown in the indicated condition under constant shaking and 30°c in 96-well plates in an automated handling robot (EVOware, Tecan Inc.). OD was measured automatically every ~30min for 65 hours using infinite200 reader.

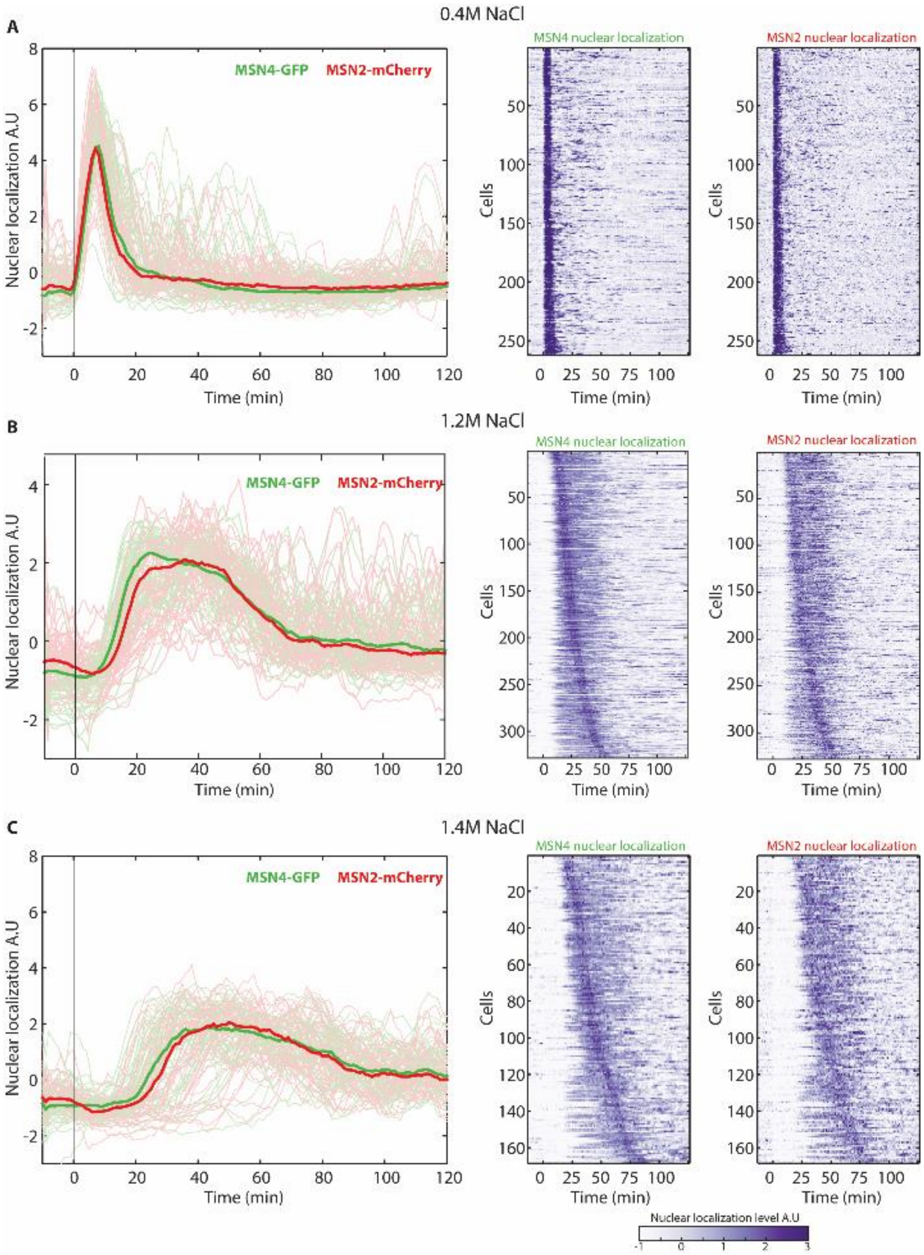


**Fig. S2.** Nuclear translocation of Msn2,4: Single cells expressing Msn4-GFP and Msn2-mChrerry fusion proteins were tracked using microfluidics-coupled live microscopy in 0.4/1.2/1.4 M NaCl. (left) Localization dynamics following exposure to stress is shown as the medians, and the single cell traces as shaded lines. (right) Individual nuclear localization traces of both Msn2 and Msn4 are shown, with cells in both columns presented in the same order.

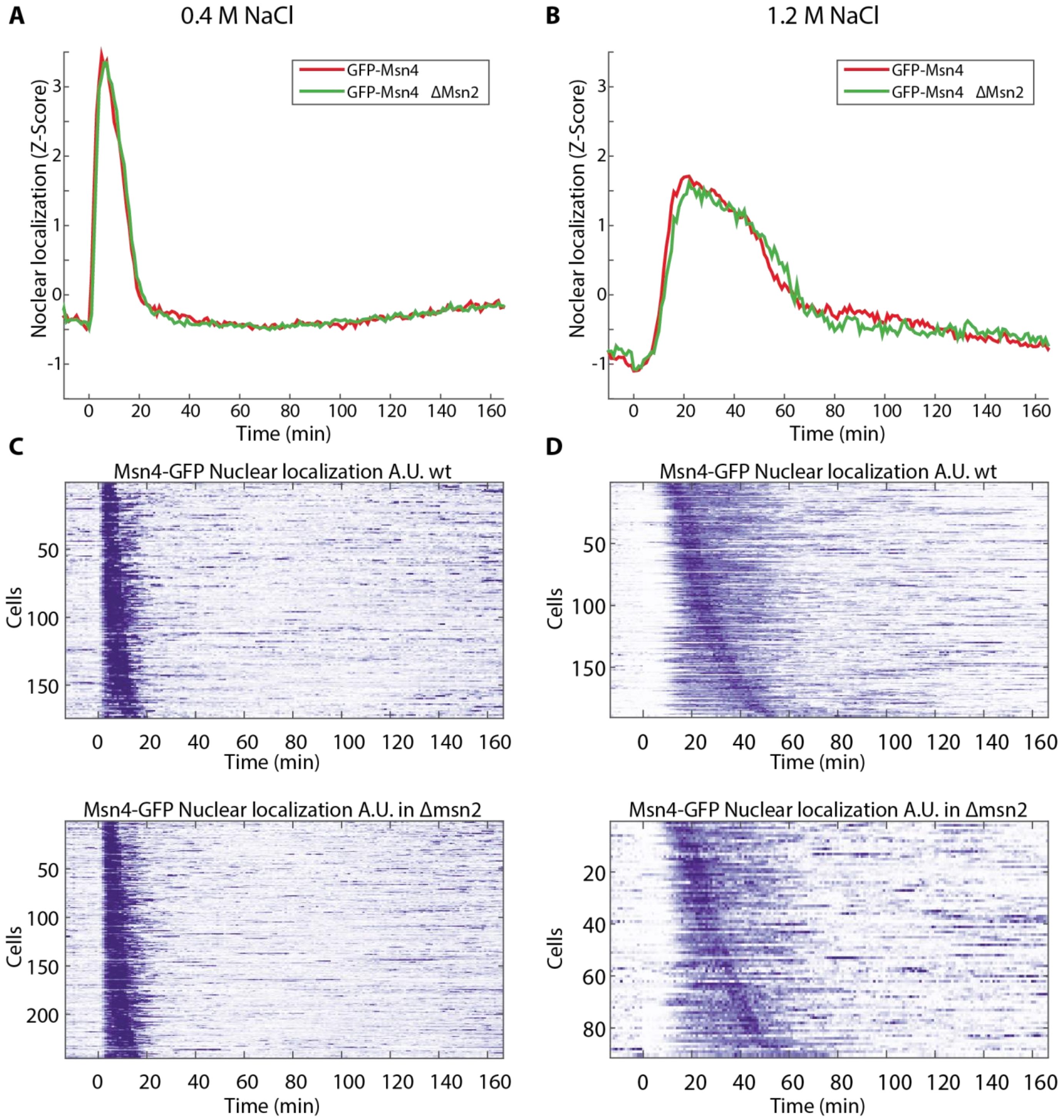


**Fig. S3.** Nuclear translocation of Msn4 in wild-type cells and cells that are deleted of msn2. (A,B) Localization dynamics following exposure to 0.4/1.2M NaCl is shown as the median. (C,D) Individual traces of Msn4 in wt cells of cells deleted of msn2.

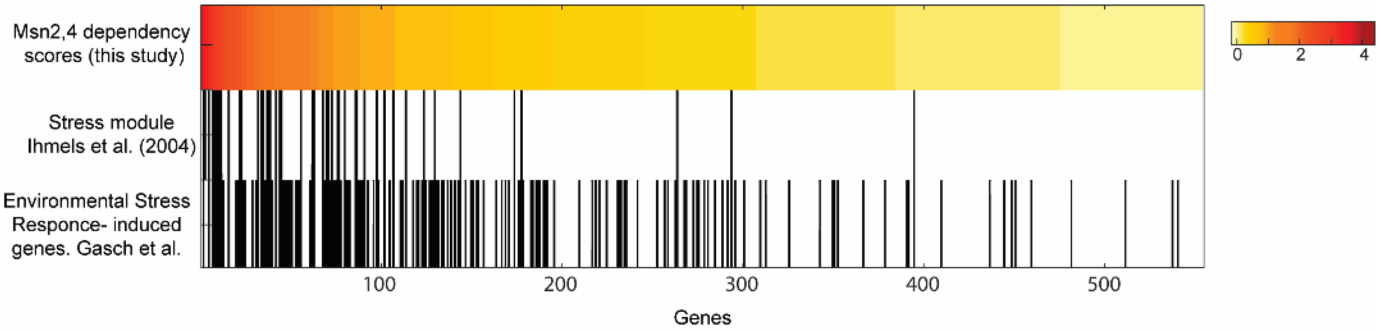


**Fig. S4.** Msn2,4 dependent genes. We calculated Msn2,4 dependency score for each gene as the average over all conditions, of ratio between wild-type induction and the double msn2,msn4 deletion strain induction. 500 top Msn2,4 dependent genes are ordered by this score. Shown are the scores and an indication if the genes are part of the written published datasets[12,64] (black: gene is part of the group, white: gene is not part of the group.).

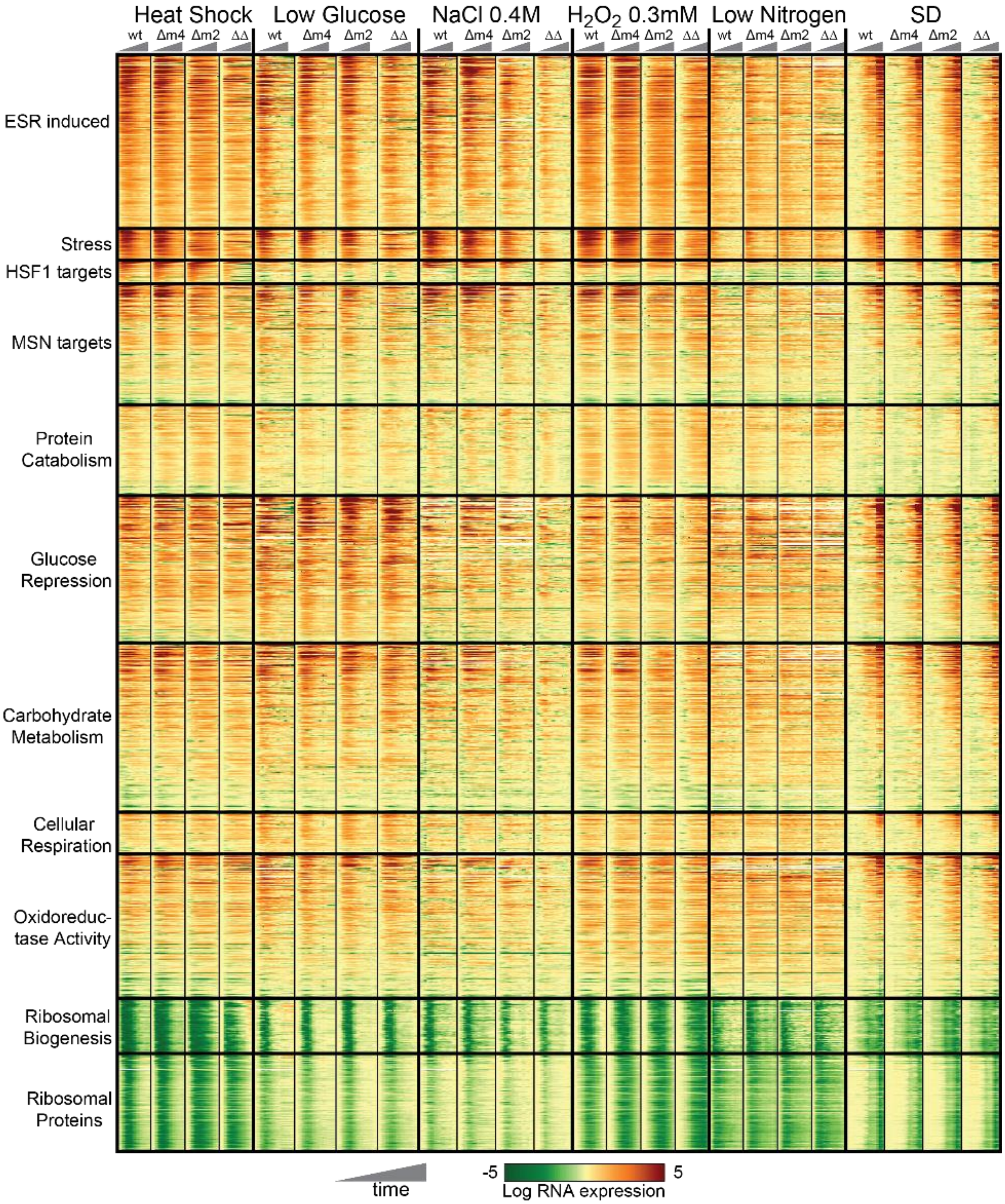


**Fig. S5.** *RNA expression in all Conditions.*

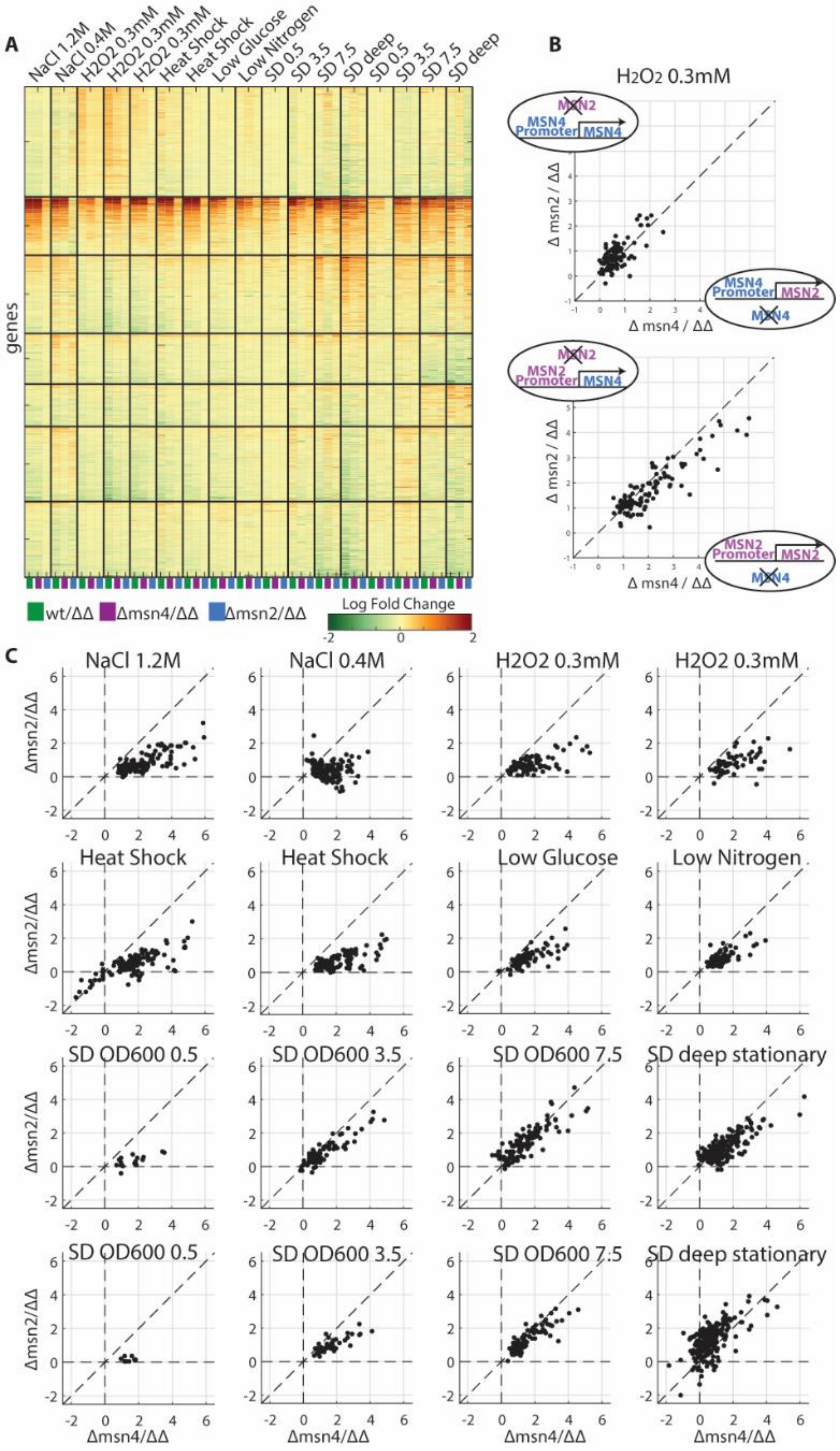
A
NaCl 1.2M
NaCl 0.4M
H2O2 0.3mM
H2O2 0.3mM
H2O2 0.3mM
Heat Shock
Heat Shock
Low Glucose
Low Nitrogen
SD 0.5
SD 3.5
SD 7.5
SD deep
SD 0.5
SD 3.5
SD 7.5
SD deep
genes
wt/ΔΔ
Δmsn4/ΔΔ
Δmsn2/ΔΔ
Log Fold Change
B
H2O2 0.3mM
MSN2
MSN4 Promoter
MSN4
Δ msn2 / ΔΔ
Δ msn4 / ΔΔ
MSN4 Promoter
MSN2
MSN4
MSN2 Promoter
MSN2
C
NaCl 1.2M
NaCl 0.4M
H2O2 0.3mM
Heat Shock
Low Glucose
Low Nitrogen
SD OD600 0.5
SD OD600 3.5
SD OD600 7.5
SD deep stationary
Δmsn2/ΔΔ
Δmsn4/ΔΔ

**Fig. S6.** *Msn2 and Msn4 induce the same target genes*. (A) Clustering of all genes in all the conditions and repeats that we checked. For each experiment of the stress perturbations we calculated for each strain the A.U.C, and for cells growing into stationary phase we used expression in different ODs. We then calculated the fold change of wt or single deletions to the double deletion strain, and used these values to cluster genes. (B) Swapping Msn2,4 promoter. Shown is the fold change of single deletions to double deletions, in the indicated strains. (C) Each dot represent a gene that was >2 fold higher in the wt then the double deletion. Shown are fold changes of single deletions to double deletion.

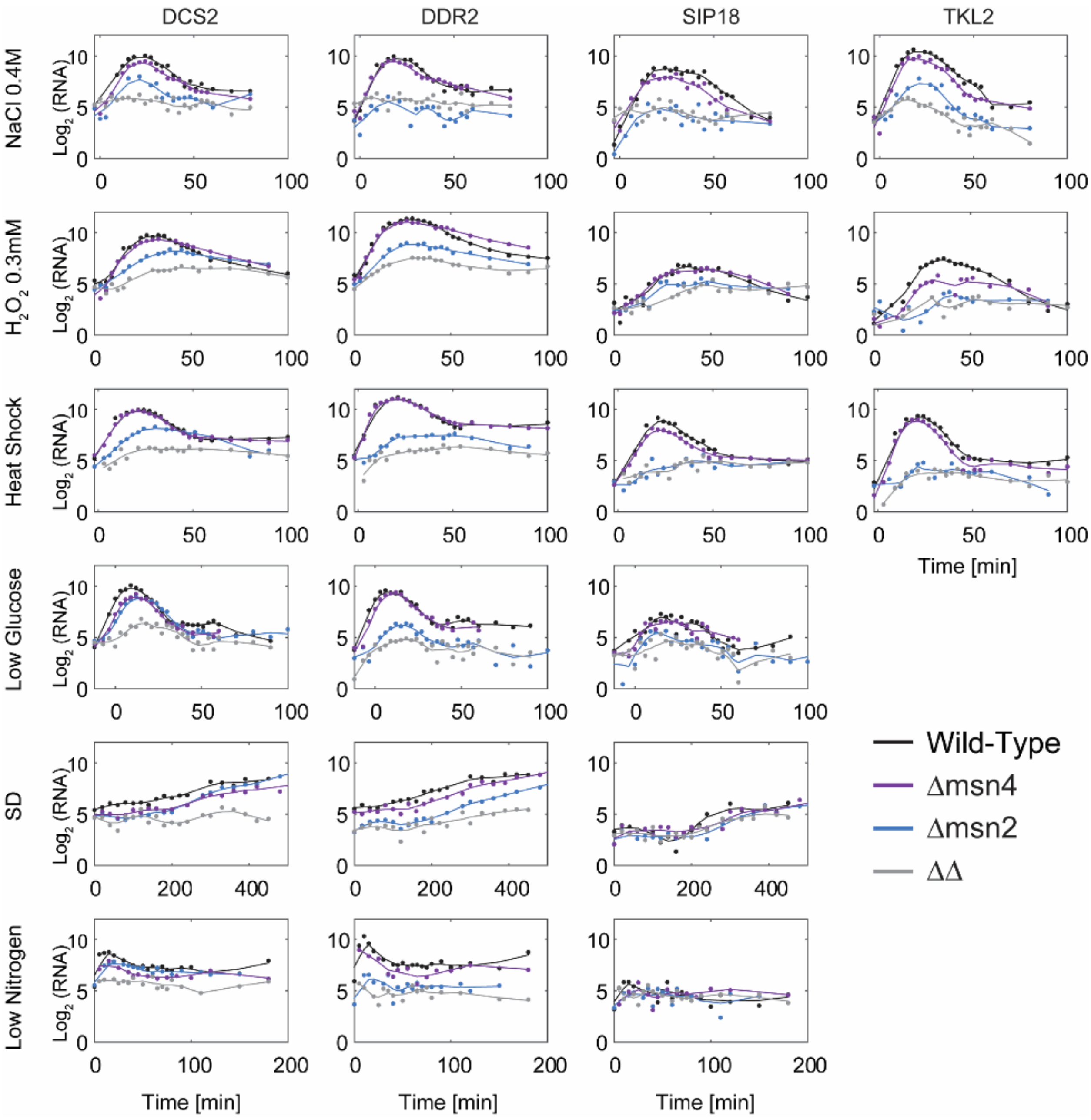


**Fig. S7.** *Response to stress of reported genes from AkhavanAghdam et al*[39]. Plotted are mRNA measurments (from this study) of the response of 4 genes reported in AkhavanAghdam et al. Shown are mRNA measurments for the wild-type, single and double deletion msn2,4 strains. Dots represents the data measurments and lines is the smoothened signal. In our high temporal resolution data there is no fundamental difference in Msn2,4 contribution to the reponse between the first two genes(DSC2,DDR2) and last two genes (SIP18,TKL2) as was reported. In all of these gense Msn4 deleted strains show similar epression and dynamics to the wt strain, where Msn2 deleted stains reduces the induction significantly.

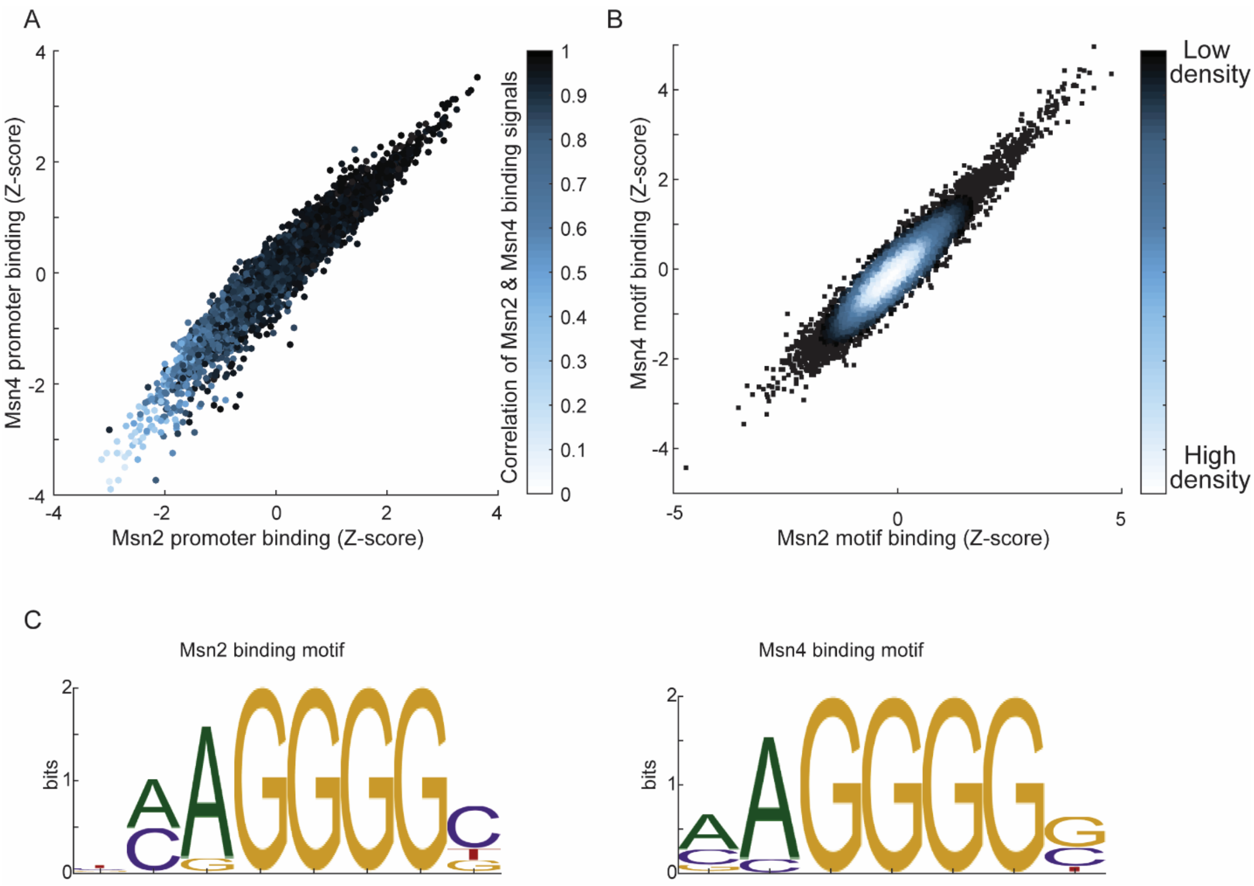


**Fig. S8.** *Msn2,4 prefer the same binding sequence and the same promotors in-vivo.* (A) Msn2 and Msn4 binding to all the promotors. Sum of the normalized CHeC-seq signal of each factor was calculated for all the promotes, in >4 repeats. Shown is a the z-score of the median of all repeats. Color represents the correlation of Msn2 and Msn4 binding signal on the promoters. (B) Density plot comparing Msn2 and Msn4 *in-vitro* binding to all possible (8192) 7-DNA base-pair sequences. For each 7-mer, the mean signal of all its appearances in all promoters was calculated for Msn2 and Msn4. Shown is the density plot of the z-scores of all possible 7-mers. (C) Motifs found in our data for Msn2 and MSN4.

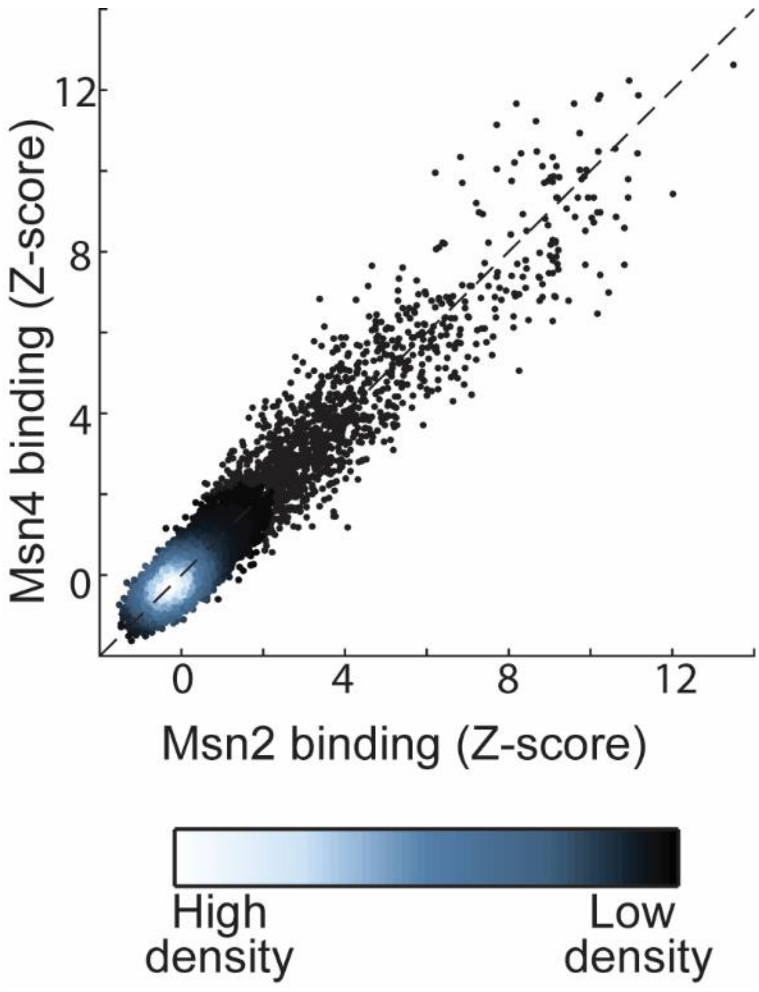


**Fig. S9.** *Msn2,4 prefer the same binding sequence site in-vitro.* Density plot comparing Msn2,4 *in-vitro* binding to all possible (32,896) 8-DNA base-pair sequences. Data from Siggers et al.[41]

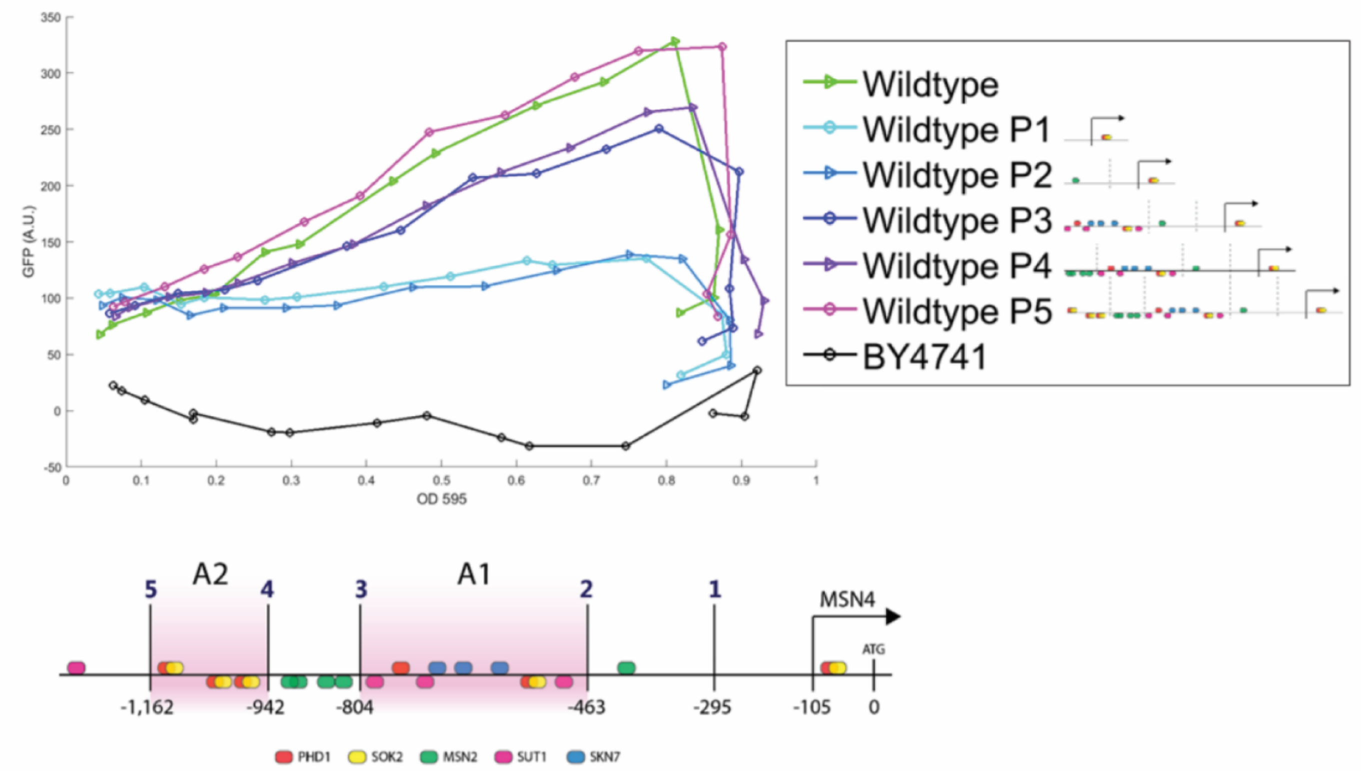


**Fig. S10.** *Msn4 promoter regions*. Median expression of Msn4-GFP along the growth curve in strains with full and partial *MSN4* promoter. The scheme shows the promoter regions that induce Msn4 at high ODs.

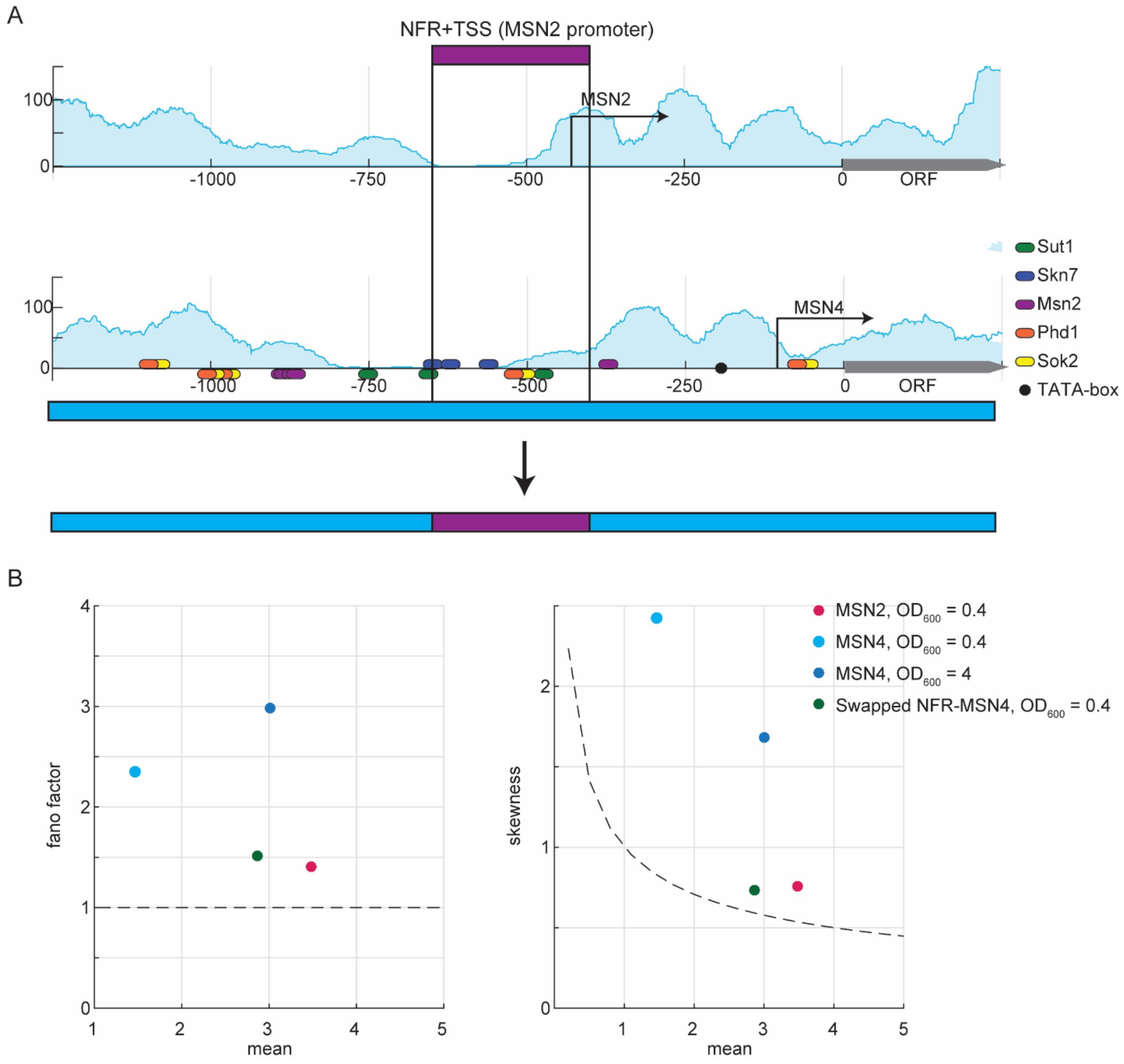


**Fig. S11.** *Msn2 NFR and TSS promoter region determine the expression level and noise.* (A) A scheme of the strain we used – *MSN4* promoter with a swap with *MSN2 NFR+TSS* in the same position. (B) smFISH results of the swapped strain and the wt Msn2,4 in the indicated ODs.

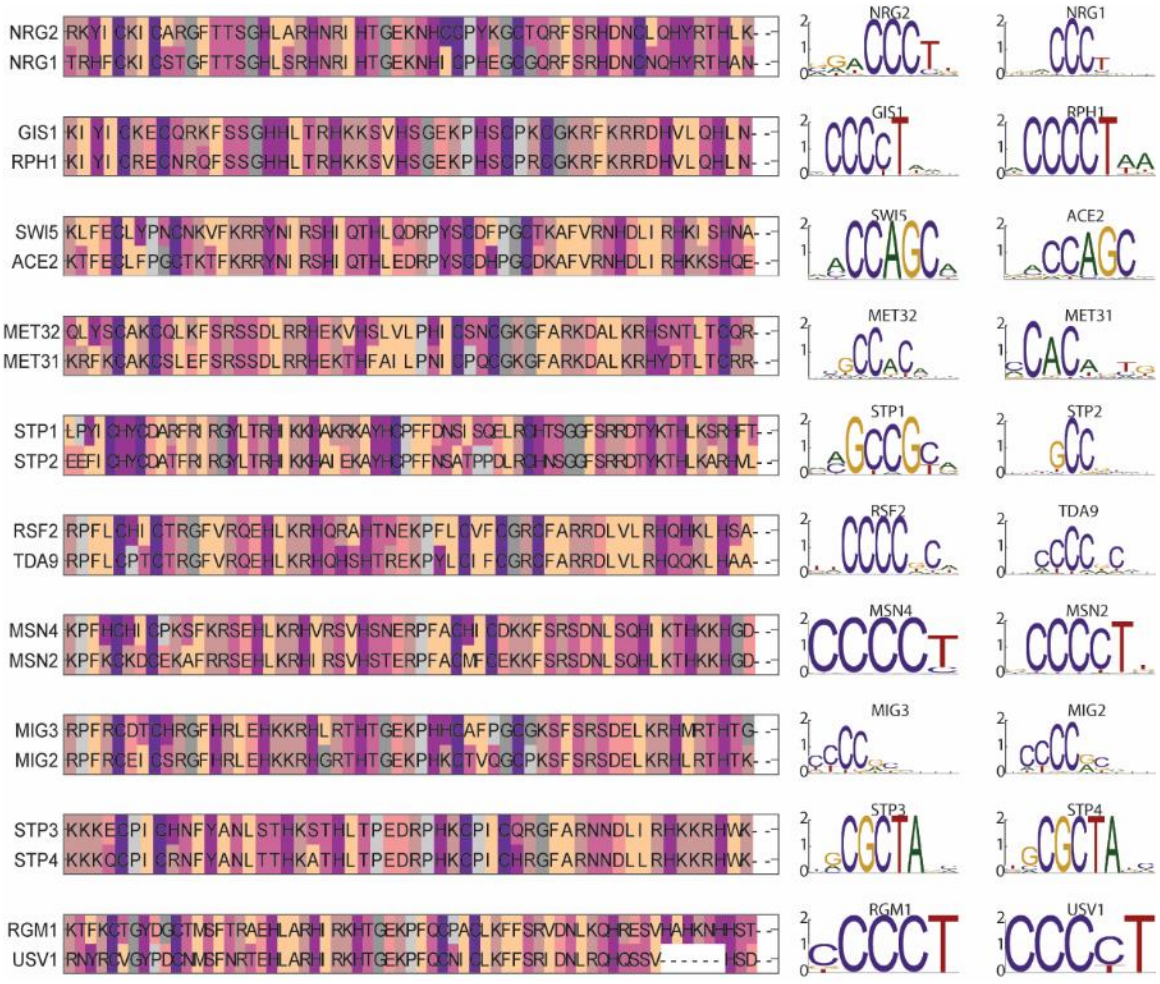


**Fig. S12.** *All S. cerevisiae Zinc finger TFs duplicates from WGD.* (left) Alignment of binding domains of all duplicated pairs. (right) DNA binding motifs of the pairs from YeTFaSCo[65].

**Table S1.** Yeast strains used in this study.

| | Strain | Genotype | Source |
|---|---|---|---|
| 1 | BY4741 | MATa his3Δ1 leu2Δ0 met15Δ0 ura3Δ0 | - |
| 2 | BY4742 | MATα his3Δ1 leu2Δ0 lys2Δ0 ura3Δ0 | - |
| 3 | wild-type | BY4741 | - |
| 4 | Δmsn2 | BY4741 msn2Δ:: hphNT1 | This study |
| 5 | Δmsn4 | BY4741 msn4Δ:: natNT2 | This study |
| 6 | Δmsn2Δmsn4 | BY4741 msn2Δ:: hphNT1 msn4Δ:: natNT2 | This study |
| 7 | Δmsn2, MSN4-promoter swap | BY4741 msn2Δ:: hphNT1 MSN4promoterΔ:: natNT2-MSN2promoter | This study |
| 8 | Δmsn4, MSN2-promoter swap | BY4741 msn4Δ:: natNT2 MSN2promoterΔ:: hphNT1-MSN4promoter | This study |
| 9 | Δmsn4, mCherry | BY4741 msn4Δ:: natNT2 HO::Kan+TDH3prom-mCherry | This study |
| 10 | Δmsn4 & MSN2-YFP | BY4741 msn4Δ:: natNT2 MSN2-YFP::kanMX4 | This study |
| 16 | Δmsn4 & MSN2 promoter library - 50 strains | BY4741 msn4Δ:: natNT2 MSN2-YFP::kanMX4 MSN2promoterΔ::URA-promoterX | This study |
| 17 | GFP-Msn4 | BY4742 GFP-MSN4 (N terminus tag), different colonies | This study, based on a strain from Yofe et al.[66] |
| 18 | GFP-Msn4 & Δmsn2 | BY4742 GFP-MSN4 msn2Δ:: hphNT1 | This study, based on strain 16 |
| 19 | GFP-Msn4 & Msn2-mCherry | BY4742 GFP-MSN4 MSN2-mCherry::His3 | This study, based on strain 16 |
| 20 | Msn2-GFP | BY4741 MSN2-GFP:: kanMX4 | This study |
| 21 | Msn2-Mnase | BY4741 MSN2-3FLAG-MNase:kanMX6 | This study |
| 22 | Msn4-Mnase | BY4741 MSN4-3FLAG-MNase:kanMX6 | This study |
| 23 | K.lactis promoter-Msn2 | BY4741 MSN2promoteΔ::K.lactisMSNpromoter | This study |
| 24 | MSN4 promoter with MSN2 promoter NFR+TSS | BY4741 MSN2promoter(400-650)Δ:: MSN4promoter(400-650) | This study |

**Table S2.** MSN2 smFISH probes, CAL Fluor Red 590

| PROBE # | PROBE (5'-> 3' | PROBE # | PROBE (5'-> 3' | PROBE # | PROBE (5'-> 3' |
|---|---|---|---|---|---|
| 1 | tgaaatcatggtcgaccgtc | 17 | agcatggagtctatgttcag | 33 | ccactttcgcaataacggac |
| 2 | ctactcatgctttctatggg | 18 | gctaaatcttcggcgtgata | 34 | tcattactcaagcctgtagt |
| 3 | ttgggttattattctccacg | 19 | tggttccaaaggtccaaaga | 35 | ttcaccttcctctgtcaaaa |
| 4 | ccgcactatctaagacagtg | 20 | cttgcattggaacttgaggt | 36 | ggagccatattcatttgagt |
| 5 | ccccaaattcagtgaagttt | 21 | ttgcgttcatagtagtggta | 37 | ttgcaagagacgtggaggat |
| 6 | gtagtggatggtatcgtttc | 22 | gttgccagcaatatttgagt | 38 | tccgaagggaagaaccgatt |
| 7 | tttgttgctatcagctttca | 23 | ctatggtagcgtcattgttt | 39 | tcaaaggcacagcagacttc |
| 8 | tcaatagttcttgcatcgcg | 24 | agtggatgtgcccaaattat | 40 | tgttgatttgtaagcggcac |
| 9 | gagccactagcattattagt | 25 | ggcaagcagatttgttcttg | 41 | gagttgacactactactgct |
| 10 | gagttgtgtattgatttgcc | 26 | atctttgtcatagcattggc | 42 | tgtcattgattttctcctgt |
| 11 | gtgacggtgattgtaacgga | 27 | tgctgtaattgctgacttgg | 43 | cgacacttgatcttctggac |
| 12 | ggattttgaggtaccgatga | 28 | tgtgttgaactcggtttagc | 44 | tcgagttcctttgttgattc |
| 13 | cttgctgtatttatgggagg | 29 | ttatgtgacgaggtgagctg | 45 | tgtgacagtggaacggtttc |
| 14 | ccatttgaagtttgaggcga | 30 | aaagaagctctttgccttct | 46 | atgccttttcaaatgttcgc |
| 15 | gcaccattaggtaaggacag | 31 | tgcttattaaccaggtcgta | 47 | ggtcgttcgttagagtgaac |
| 16 | acccgtagaattggagtttg | 32 | aattcggcagcatatcgttc | 48 | gtcttgatgtgttgcgacaa |

**Table S3.** MSN4 smFISH probes, CAL Fluor Red 590

| PROBE # | PROBE (5'-> 3' | PROBE # | PROBE (5'-> 3' | PROBE # | PROBE (5'-> 3' |
|---|---|---|---|---|---|
| 1 | ttaggtccgaagactagcat | 17 | catgttggtgtggcgataat | 33 | ggttgcatctaagctattgt |
| 2 | ttgtttcttgtttgcgtgac | 18 | aggtttgtcttgggagaagt | 34 | gggtctacgttattgtcgaa |
| 3 | ggctcgttcattatagacga | 19 | atattgaaggttctgcctca | 35 | taaactgttgatcctgagcc |
| 4 | cagaaacgttggttgtgctc | 20 | gtttggtaacctagcatcta | 36 | cagaggcattatccgacaac |
| 5 | gctattgttcgcagaattgt | 21 | cgttcaatcctgttgtactg | 37 | accagaagtcgccaatttag |
| 6 | aaggtatattccggcgaaga | 22 | cggcatgaacttcaatgctt | 38 | tttggaaagctggtgtaggt |
| 7 | ctatgcggagaatccattga | 23 | aaggaagaggtaaccacctc | 39 | tggtgagcttgactagtagg |
| 8 | attggagtattagtggcgtc | 24 | gttgagtgggtttactgttg | 40 | gttgttgatgatgttgagct |
| 9 | cgtattagttgtcgctgtta | 25 | gcagtaccttcagggataat | 41 | gacgactttcttcttcttgt |
| 10 | ggcggtatcctttaaattca | 26 | ggtgaaatggttgatccacg | 42 | aaccttaccattgttgttgt |
| 11 | gaatttttccacttccagtt | 27 | gctcagagcttgagaaacgt | 43 | atagatttccttttccgagg |
| 12 | cattcacgaactgtgcgtta | 28 | tttgttataccagagtcctt | 44 | tttatcgtagttgttggggt |
| 13 | agcattactgctttgtttct | 29 | gctatgggagatgcgattta | 45 | cttctcacagtctttacact |
| 14 | cttatcaccaccatttttct | 30 | ttaagggcaaagaggcacgc | 46 | cggatcttatatgccttttc |
| 15 | gggtggctactgaactatat | 31 | gcggaatcagacagaggatt | 47 | tacaagcaaaagggcgttcc |
| 16 | ctagttgctccaagtttttc | 32 | caccaaaagcatcgtcttca | 48 | cgtgctttttgtgagttttt |
